# Repurposed Drugs Block Toxin-Driven Platelet Clearance by the Hepatic Ashwell-Morell Receptor to Clear *Staphylococcus aureus* Bacteremia

**DOI:** 10.1101/2020.07.06.190322

**Authors:** Josh Sun, Satoshi Uchiyama, Joshua Olson, Yosuke Morodomi, Ingrid Cornax, Nao Ando, Yohei Kohno, May M. T. Kyaw, Bernice Aguilar, Nina M. Haste, Sachiko Kanaji, Taisuke Kanaji, Warren E. Rose, George Sakoulas, Jamey D. Marth, Victor Nizet

**Author notes:** These authors contributed equally (Josh Sun, Satoshi Uchiyama). Correspondence: Victor Nizet.

## Abstract

*Staphylococcus aureus* (SA) bloodstream infections cause high morbidity and mortality (20-30%) despite modern supportive care. In a human bacteremia cohort, development of thrombocytopenia was correlated to increased mortality and increased α-toxin expression by the pathogen. Platelet-derived antibacterial peptides are important in bloodstream defense against SA, but α-toxin decreased platelet viability, induced platelet sialidase to cause desialylation of platelet glycoproteins, and accelerated platelet clearance by the hepatic Ashwell-Morell receptor (AMR). Ticagrelor (Brilinta^®^), a commonly prescribed P2Y12 receptor inhibitor used post-myocardial infarction, blocked α-toxin-mediated platelet injury and resulting thrombocytopenia, thus providing protection from lethal SA infection in a murine intravenous challenge model. Genetic deletion or pharmacological inhibition of AMR stabilized platelet counts and enhanced resistance to SA infection, and the anti-influenza sialidase inhibitor oseltamivir (Tamiflu^®^) provided similar therapeutic benefit. Thus a “toxin-platelet-AMR” regulatory pathway plays a critical role in the pathogenesis of SA bloodstream infection, and its elucidation provides proof-of-concept for repurposing two FDA-approved drugs as adjunctive therapies to improve patient outcomes.

## INTRODUCTION

*Staphylococcus aureus* (SA) is one of the most important human bacterial pathogens, as the second leading cause of bloodstream infections (bacteremia) and the leading cause of infective endocarditis *(1)*. Despite modern supportive measures, overall mortality in SA bacteremia has not declined in decades and remains unacceptably high (20-30%), with significant risk of complications including sepsis syndrome, endocarditis and metastatic foci of infection (e.g. osteomyelitis) *(2)*. High risk populations include the elderly, diabetics, surgical and hemodialysis patients *(3)*. Multidrug resistance (e.g. methicillin-resistant SA, MRSA) is prevalent and associated with adverse outcome and increased medical costs *(4)*.

The high incidence of SA bacteremia signifies a remarkable capacity of the organism to resist host innate defense mechanisms that function to prevent pathogen bloodstream dissemination *(5)*. Extensive research has focused on SA virulence factors that counteract opsonization by serum complement *(6)*, surface-anchored protein A that impairs Fc function of antibodies *(7)*, and the pathogen’s numerous resistance mechanisms to avoid phagocytosis and oxidative burst killing by neutrophils *(8)*.

Comparatively less is understood about how SA interacts with circulating platelets. These abundant, small anucleate cells are best known for their central role in hemostasis, but increasingly appreciated to possess bioactivities relevant to immune defense *(9)*. Platelets can act as mechano-scavengers to bundle bacteria *(10)* and enhance the function of professional phagocytic cell types such as neutrophils *(11)*, macrophages *(12)* and hepatic Kuppfer cells *(13, 14)*. Platelets express several Toll-like receptors that recognize pathogen-associated molecular patterns *(15)* to activate their release of pro-inflammatory cytokines (e.g. interleukin-1β) *(16)* and antimicrobial peptides including platelet microbicidal protein (tPMP) and human beta-defensin-1 (hBD-1) with direct antibacterial actions *(12, 17, 18)*. SA activates platelets via integrin GP11b/IIIa, FcγRIIa receptor and ADAM-10-dependent pathways *(19–21)*, and the pathogen induces platelet aggregation via its “clumping factors” ClfA and ClfB *(22, 23)*.

In a cohort of 49 patients with SA bacteremia, we report a strong association of mortality with lowered platelet count (thrombocytopenia) rather than changes in leukocyte count. Of note, platelet depletion in mice was recently shown to impair SA clearance *(12, 24)*. Our mechanistic analysis with human platelets *ex vivo* and murine platelets *in vivo* revealed a critical activity of platelets in direct killing of SA. The pathogen attempts to counteract this defense by deploying a pore-forming toxin (α-toxin) to (i) disrupt platelet antimicrobial activity and (ii) accelerate sialidase-dependent platelet clearance through the hepatic Ashwell-Morell receptor (AMR). Elucidation of this “toxin-platelet-AMR” regulatory pathway guided us to therapeutic repurposing of two FDA-approved drugs to preserve platelet homeostasis, thereby providing significant host protection in experimental SA bloodstream infection.

## RESULTS

### Platelets are essential for blood immunity against SA bacteremia

The normal human platelet count ranges from 150,000-450,000/mm^3^ of blood. In 49 consecutive patients with SA bacteremia (blood culture growing MRSA or MSSA) identified prospectively at an academic medical center in Madison, WI *(25)*, we observed a strong association of patient mortality with thrombocytopenia (platelet count <100,000/mm^3^) on the initial blood sample and not abnormally elevated or reduced leukocyte count (**Fig. 1, A and B**). Indeed, two patients with thrombocytopenia failed to clear their bacteremia despite antibiotic therapy for more than 60 days before succumbing. No significant correlation was observed between platelet count and serum levels of pro-inflammatory cytokine IL-1β nor APACHE score, a clinical metric of disease severity (**Fig. 1A**). These data are consistent with a prior single center study in Israel showing thrombocytopenia (and not leukocyte count) to be a significant risk factor for 30-day all-cause mortality in SA bacteremia, although overall mortality rates were much higher than other published series (56.2% in thrombocytopenic group vs. 34% with normal platelets counts) *(26)*. These clinical studies suggest that circulating platelets, and not white blood cells, could play the dominant role in resolution of SA during bloodstream infection. Indeed, when directly compared, the second most common Gram-positive bacterial pathogen associated with human bacteremia, *Streptococcus pneumoniae (1)*, SA was ~50% more resistant to killing by purified human neutrophils, but ~70% more susceptible to killing by human platelets (**Fig. 1C**).

**Fig. 1.**
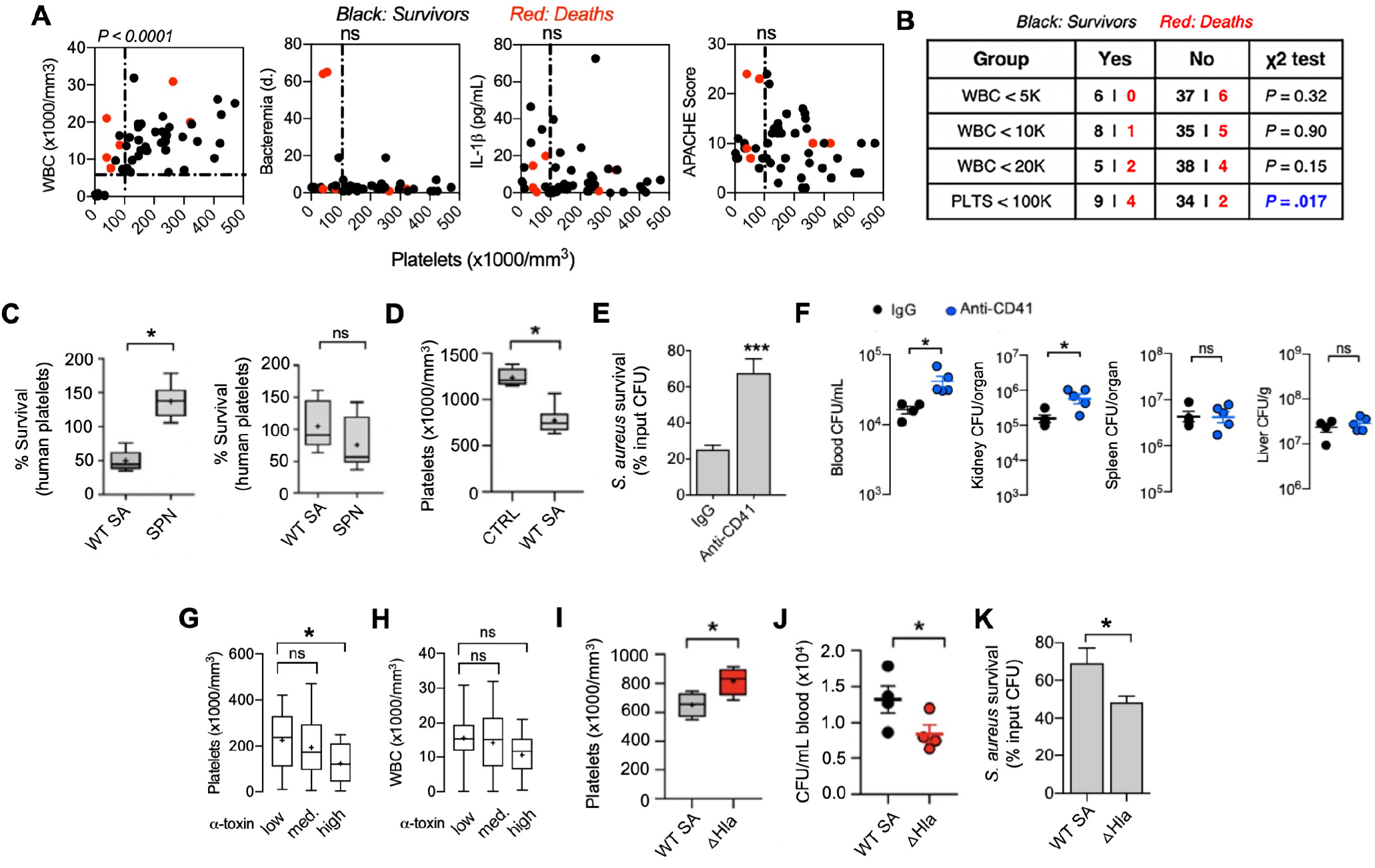
Platelets are essential for blood immunity against *Staphylococcus aureus* (SA) bacteremia and SA α-toxin induces thrombocytopenia to evade platelet-mediated microbicidal activity. **(A)** Correlation of circulating platelet counts with leukocyte counts and duration of bacteremia in 49 consecutive patients with SA bacteremia from a tertiary medical center; Spearman’s rank correlation coefficient compared variables. **(B)** Mortality in patient cohort associated with different leukocyte and platelet count cutoffs; Chi-square without Yates correction, P < 0.05 considered significant. **(C)** Washed isolated human platelets and neutrophils exposed to SA or SPN at MOI = 0.01 for 2 h. Samples sonicated, serial diluted, and plated on THA plates for enumeration of bacterial colony forming units (CFU). **(D)** Reduction in platelet count 2 h post intravenous infection of mice with SA (n = 8) vs. non-infected littermate control (n = 4). Experiment reproduced x 2 and data pooled; data represented as mean ± SEM. **(E)** *Ex vivo* killing of SA upon 2 h co-incubation with blood collected from mice 16 h after treatment with anti-CD41 antibody (n= 9) or IgG control (n= 12). **(F)** Mice treated with platelet-depleting anti-CD41 antibody (n = 5) or IgG control (n = 4) for 16 h prior to intravenous SA infection. Organs harvested and CFU enumerated 2 h post-infection in triplicate for each sample. **(G)** Assessment of α-toxin production by the infecting SA isolate in 49 consecutive bacteremia cases and its association with patient platelet counts and **(H)** white blood cell counts. **(I)** Wild-type SA (n = 4) or isogenic ΔHla mutant (n = 4) intravenously challenged outbred CD-1 mice. Blood harvested by cardiac puncture and CBC obtained 4 h post-infection. **(J)** Wild-type SA (n = 4) or isogenic ΔHla mutant (n = 4) intravenously challenged outbred CD-1 mice. Blood harvested by cardiac puncture and enumeration of colony forming units (CFU) 4 hours post-intravenous infection. **(K)** *Ex vivo* killing of SA by freshly isolated human platelets (2 h co-incubation) vs. isogenic ΔHla mutant. All data represented as mean ± SEM and representative of at least 3 independent experiments. Statistical significance was determined by unpaired two-tailed Student’s T-test (C-F, I, J, K) or one way analysis of variance (ANOVA) with Bonferroni’s multiple comparisons test (G,H). *P < 0.05. For floating bar graphs, + denotes the mean, whiskers represent min. to max, and floating box represents 25th to 75th percentile.

We pursued this association further using an *in vivo* model of SA bacteremia (MRSA strain USA300 TCH1516) established by intravenous (i.v.) tail vein injection in mice, where normal platelet count ranges from 900,000 to 1.4 million cells/mm^3^ blood *(27)*. Within 2 h of SA challenge, the circulating platelet count of infected mice was reduced by ~40% from baseline levels (1237 ± 49.82 vs. 772 ± 49.91) (**Fig. 1D**). To verify that reduced platelet count was indicative of a functional immune deficiency, we used an anti-CD41 antibody to deplete mice of platelets to 17% of baseline levels (**fig. S1**). The drawn blood of the thrombocytopenic animals was impaired in *ex vivo* killing of SA (67.5 ± 7.9% surviving CFU vs. 25 ± 2.1% surviving CFU in normal blood) (**Fig. 1E**), and bacterial burdens in blood and kidneys of thrombocytopenic mice were significantly increased vs. untreated mice within 2 h following i.v. SA challenge (**Fig. 1F**).

### SA α-toxin induces thrombocytopenia to evade platelet-mediated microbicidal activity

A major SA secreted virulence factor, the pore-forming α-toxin (Hla), induces platelet cytotoxicity and aberrant aggregation after binding its protein receptor A-disintegrin metalloprotease-10 (ADAM-10) on the platelet surface *(14, 20, 28)*. Using ImageJ densitometric analysis of anti-Hla western immunoblots, we grouped the SA bacteremia isolates from our clinical cohort into low- (n= 18), medium- (n=15), and high- (n=16) α-toxin producers (**fig. S2**). A significant correlation was seen between high-level α-toxin production and thrombocytopenia (**Fig. 1G**), compared to the low-level and medium-level α-toxin producing groups. There was no significant correlation between α-toxin production and leukocyte counts (**Fig. 1H**) For comparison to the wild-type (WT) parent SA strain in analyses of platelet interactions, we constructed a Δ*hla* knockout strain by precise allelic replacement (**fig. S3**). In the mouse i.v. challenge model, the SA Δ*hla* mutant induced less thrombocytopenia (**Fig. 1I**) and was more rapidly cleared from the blood circulation (**Fig. 1J**) than the WT parent strain. *Ex vivo*, the SA Δ*hla* mutant was more susceptible to killing by purified human platelets (**Fig. 1K**). Together these studies indicate that the virulence effects of SA α-toxin extend to evasion of direct platelet-mediated antibacterial killing.

### FDA-approved P2Y12 inhibitor ticagrelor blocks SA α-toxin-mediated platelet cytotoxicity

Inhibition of platelet activation is the target of antithrombotic drug therapy designed to reduce the risk of cardiovascular death, myocardial infarction (MI), and stroke in patients with acute coronary syndrome or a history of MI, beginning with classical studies of aspirin (acetylsalicylic acid, ASA) in the 1970s, then extending to newer selective inhibitors of adenosine signaling through the platelet P2Y12 receptor (clopidogrel, prasugrel, ticagrelor) *(29)*. However, the effect of “antiplatelet” drugs on the direct antibacterial properties of platelets has not been reported. Of potential relevance, a clinical study of 224 consecutive patients with community-acquired pneumonia found that those receiving antiplatelet therapy (ASA and/or clopidogrel) for secondary prevention of cardiovascular disease had reduced need for intensive care unit treatment (odds ratio 0.19, 95% confidence interval 0.04−0.87) and shorter hospital stays (13.9 ± 6.2 vs. 18.2 ± 10.2 days) compared to their age-matched cohort *(30)*. Additional human retrospective or matched cohort studies of endocarditis, bacteremia or sepsis (not restricted to SA) have provided similar hints of improved clinical outcome among patients receiving antiplatelet drugs *(31–34)*.

To discriminate the effect of the two antiplatelet drug classes on SA killing, we pretreated freshly isolated human platelets for 15 min with ASA or ticagrelor (chosen since clopidogrel is a prodrug requiring hepatic conversion *in vi*vo) and co-incubated them with the bacteria. Within 2 h, ticagrelor-treated platelets showed a 2.2-fold enhancement in SA killing vs. untreated controls (**Fig. 2A**), whereas ASA did not significantly alter platelet antibacterial activity. In contrast, ticagrelor did not promote macrophage or neutrophil killing of SA, did not alter neutrophil extracellular trap production, and did not directly inhibit SA growth (**fig. S4, A-D**). Upon direct co-incubation of SA with human platelets in a tissue culture well, severe platelet damage was evident by transmission electron microscopy; however, ticagrelor treatment preserved platelet structural integrity against SA-induced injury (**Fig. 2B**). As α-toxin is the principal driver of SA platelet toxicity, we measured α-toxin-induced lactate dehydrogenase (LDH) release from platelets treated with ticagrelor, ASA (COX-1 inhibitor), or specific small molecule inhibitors of other known platelet activation receptors (CD41, PAR-1, PAR4). Among these agents, only ticagrelor significantly inhibited SA α-toxin-induced platelet LDH release (**Fig. 2C**), doing so in a dose-dependent manner (**Fig. 2D**). The deleterious effect of α-toxin on platelets involves activation of its receptor protease ADAM-10 leading to intracellular Ca2+ mobilization *(20)*, and these biological effects were both significantly inhibited by ticagrelor as measured in specific assays (**Fig. 2, E and F**). While ticagrelor did not alter the amount of ADAM-10 expressed on the platelet surface (**Fig. 2G**), the P2Y12 inhibitor drug blocked SA-induced ADAM-10-dependent shedding of platelet glycoprotein-6 (GP6) (**Fig. 2H**). SA exposure also induced P-selectin, a transmembrane protein specific to alpha granules that is translocated to the platelet surface upon activation (**Fig. 2I**), and raised surface levels of CD63, a marker of dense granule mobilization (**fig. S5A**). The SA-induced upregulation of P-selectin and CD63 were both blocked upon ticagrelor treatment (**Fig. 2I, fig. S5A**). Neither SA nor ticagrelor significantly affected platelet beta-galactosidase activity, another lysosomal marker (**fig. S5B**).

**Fig. 2.**
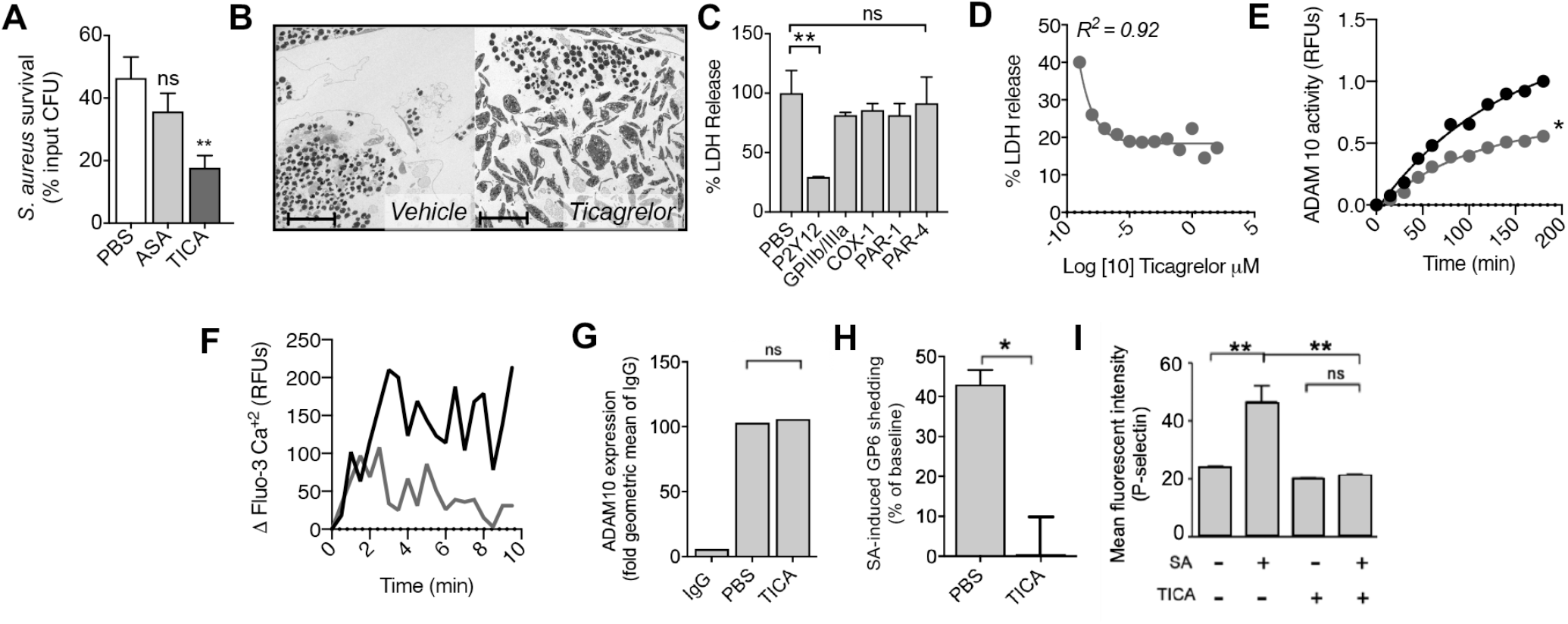
FDA-approved P2Y12 inhibitor ticagrelor blocks SA α-toxin-mediated platelet cytotoxicity. **(A)** Effect of 10 μM aspirin (ASA) and 10 μM ticagrelor (TICA) (15 min pretreatment *ex vivo*) on human platelet killing of MRSA for 2 h (n = 9). Experiments performed in triplicate and repeated three times. **(B)** Representative transmission electron microscopy image of platelets pre-treated with or without 10 μM TICA and exposed to MRSA at MOI: 0.1 for 2 h. **(C)** P2Y12 inhibitor (TICA) pretreatment blocks human platelet cytotoxicity by 5 μg/ml purified α-toxin as measured by LDH release (n = 3) in a **(D)** dose-dependent manner. Inhibitors: P2Y12 (ticagrelor), GPIIb/IIIa (eptifibatide), COX-1 (SC560), PAR-1 (vorapaxar), and PAR-4 (ML-354). **(E)** TICA treatment of human platelets reduces proteolytic cleavage of an ADAM10-specific fluorogenic substrate. Data representative of three independent experiments and statistical significance determined by least squares ordinary fit, *P < 0.5. **(F)** Measurement of intracellular calcium in human platelets loaded with 2 μM Fluo-3 dye and stimulated with 5 μg/mL recombinant α-toxin; calcium influx is measured every 30 sec by fluorescence and normalized to baseline (non-stimulated platelet control). (**G**) TICA treatment of human platelets did not alter surface ADAM-10 expression as determined by flow cytometry. Surface ADAM-10 expression on human platelets treated with TICA or control was measured by flow cytometry. (**H**) Human platelets with or without TICA treatment were infected with MRSA at MOI = 0.1 for 90 min. Surface glycoprotein-6 (GP6) was measured by flow cytometry and % decreased expression (GP6 shedding) calculated. (**I**) Human platelet P-selectin expression indicating platelet activation measured by flow cytometry with or without MRSA challenge (MOI = 0.1) and with or without TICA aor 90 min. All data represented as mean ± SEM and are representative of at least 3 independent experiments. Statistical significance was determined by One way ANOVA with Bonferroni’s multiple comparisons test (A,C,G), unpaired two-tailed Student’s T-test (H) and two-way analysis of variance (ANOVA) with Bonferroni’s multiple comparisons posttest (I). *P< 0.05, **P < 0.005. PBS, phosphate buffered saline; ns, not significant.

### FDA-approved P2Y12 inhibitor ticagrelor protects against SA bacteremia

Inhibition of α-toxin mediated platelet cytotoxicity suggested that P2Y12 inhibition using ticagrelor could mitigate the toxin’s virulence role in driving SA-induced thrombocytopenia (**Fig. 1I**) to promote bloodstream survival of the pathogen (**Fig. 1J**). IV SA challenge in mice drove down platelet counts beginning as early as 4 h (35% decrease) and continuing through 24 h (63% decrease), with partial recovery by 72 h (35% decrease) (**fig. S6A**); bone marrow analysis at 72 h revealed increased thrombopoiesis as evidenced by greater megakaryocyte number and by ploidy distribution (**fig. S6B and C**). The rapid SA-induced thrombocytopenia was associated with platelet GP6 shedding (**fig. S6D and E**) and platelet microparticulation (**fig. S6F**), the latter determined by *in vitro* studies to be α-toxin-dependent (**fig. S6G**). Indeed, mice treated with ticagrelor maintained higher numbers of circulating platelets compared to control animals following SA i.v. infection (**Fig. 3A**), significantly reducing the bacterial burden in the blood (**Fig. 3B**) in systemic organs (kidney, liver, spleen, **Fig, 3C**), and ultimately improving survival in a 10-day mortality study (**Fig. 3D**). Blinded histological examination of tissues by a veterinary pathologist revealed the most striking treatment-associated differences in the kidneys (**Fig. 3E**) and the heart (**fig. S7**). A 4- to 10-fold reduction in bacterial micro-abscesses was identified within the renal glomeruli, tubules, and blood vessels of ticagrelor-treated mice vs. PBS control animals, corroborating the CFU quantification data. Renal microabscesses in the control group were generally larger and more densely packed with bacteria, while those in ticagrelor-treated mice were frequently disrupted by immune cell infiltrates. Together these data suggest that platelet P2Y12 inhibition blocks α-toxin and SA-mediated platelet cytotoxicity and consequent thrombocytopenia, thus enhancing the platelet-mediated clearance of the pathogen *in vitro* and *in vivo*.

**Fig. 3.**
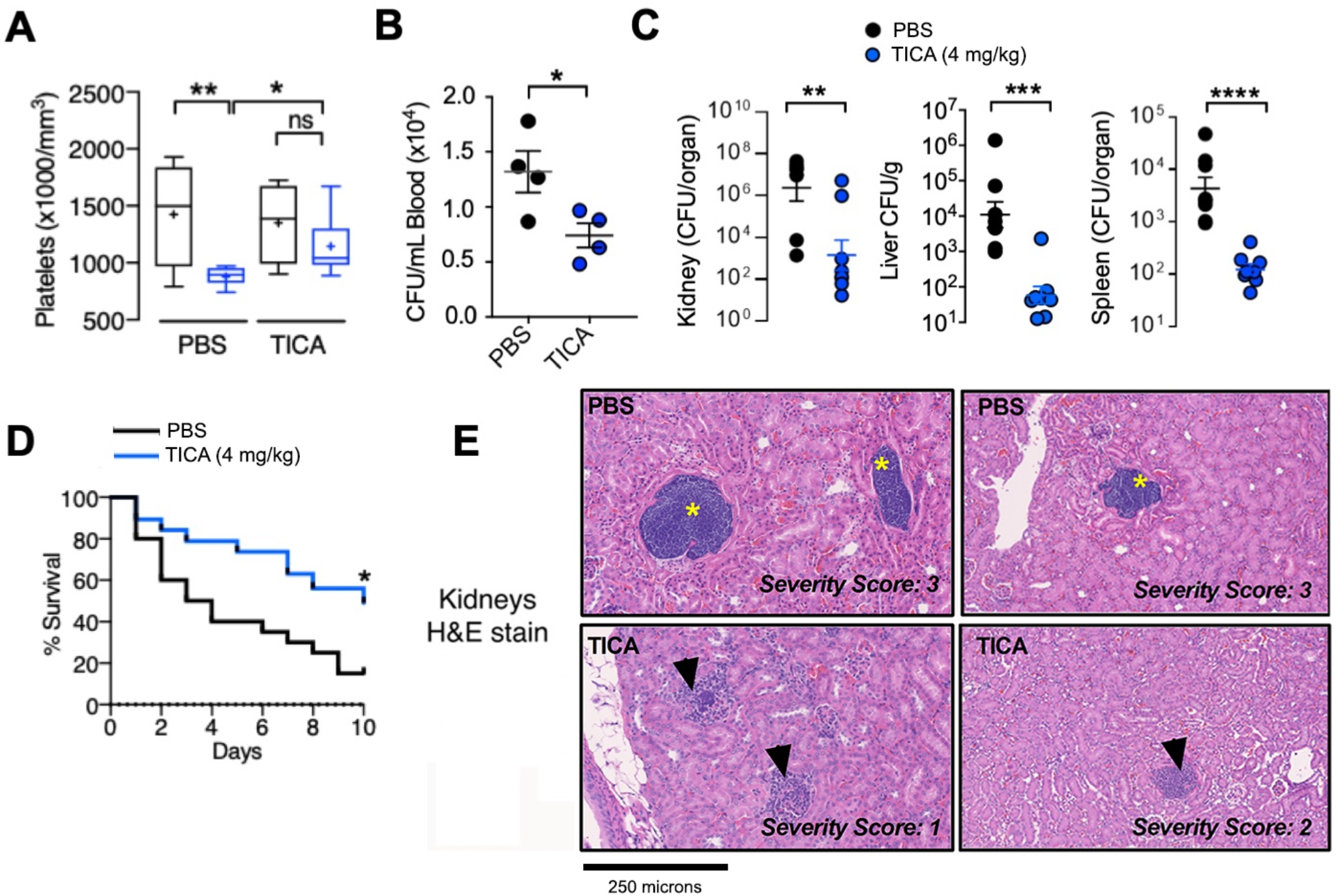
FDA-approved P2Y12 inhibitor ticagrelor protects against SA bacteremia. **(A and B)** Blood harvested by cardiac puncture and both platelets and bacterial colony forming unit (CFU) burden enumerated from outbred CD-1 mice treated with 10 μM ticagrelor (n = 9) or PBS (n = 9) control and intravenously infected with SA. **(C)** Enumeration of bacterial colony forming unit (CFU) burden at 72 h in organs of mice pretreated with vehicle (PBS) or 4 mg/kg Ticagrelor 12 h prior to intravenous SA and q 12 h thereafter; (n= 8). **(D)** Mortality curves of outbred CD-1 mice pretreated with vehicle (PBS) or 4 mg/kg Ticagrelor beginning 24 h prior to intravenous SA infection then q 12 h over a 10-day observation period (n = 20). Independent experiments were repeated x 2 and data pooled. **(E)** Hematoxylin and eosin stain (H&E) of representative histological kidney sections from mice pre-treated with PBS vehicle or 4 mg/kg ticagrelor 12 h prior to SA infection and q 12 h thereafter for 72 h; (n = 8). Yellow stars denote formation of dense bacterial colonies and black arrows represent immune infiltrate. All histological sections are representative photos of at least 6 samples per two independent experiments. Where applicable, results are represented as mean ± SEM and statistical significance was determined by unpaired two-tailed Student’s T-test (B,C), and two-way ANOVA with Bonferroni’s multiple comparisons posttest (A). For survival curves, statistical significance determined by Log-rank Mantel-Cox test (D); *P < 0.05. For floating bar graphs, + denotes the mean, whiskers represent min. to max, and floating box represents 25th to 75th percentile. Unless otherwise stated, *P < 0.05, **P < 0.005, ***P < 0.0005.

### SA α-toxin activates endogenous platelet sialidase activity

Our clinical data (**Fig. 1A**) and those of others *(26)*, coupled with our experimental work (**Figs. 1, E and F**) and prior platelet depletion studies *(12, 24)*, suggest platelet count *per se* is important in determining SA clinical outcome. Since α-toxin production correlated to thrombocytopenia in patients (**Fig. 1G**) and in experimental mouse infection (**Fig. 1I**), we hypothesized that SA deploys the toxin as a means to deplete the host of an effective circulating innate immune cell. Yet, platelet senescence and clearance are tightly regulated by multiple mechanisms, in particular the highly conserved hepatic transmembrane heterodimeric Aswell-Morell receptor (AMR) *(35)*. The AMR clears “aging” platelets with reduced terminal α2,3-linked sialic acids on their surface glycoproteins and glycolipids by engaging the exposed underlying galactose. We asked if the observed therapeutic effect of ticagrelor in SA sepsis was solely based on inhibiting platelet cytotoxicity or may further intersect with this important mechanism of platelet homeostasis.

To assess platelet sialylation state during SA bacteremia, we obtained frozen plasma from 10 randomly selected adult patients with SA bacteremia, 10 patients with *Escherichia coli* bacteremia, and 5 healthy controls. We found there was a significant increase in exposed galactose (indicative of desiaylation) on the platelets of SA-infected patients compared to the two other groups (**Fig. 4A**). However, SA lacks a bacterial sialidase (neuraminidase) present in other pathogens including *S. pneumoniae* (**Fig. 4B**). Rather, we discovered that WT SA induces sialidase activity on purified human platelets, whereas its isogenic Δ*hla* mutant derivative did not (**Fig. 4C**). An increase in platelet sialidase activity in response to WT SA or purified α-toxin, present within the platelet pellet but not released into the media, was detected in independent assays using lectin affinity and a fluorescent substrate (**fig. S8A, Fig. 4D**). Although the precise mechanism of its transfer is not established, Neu1 is the main endogenous sialidase that translocates from lysosomal stores to the platelet surface to target glycoproteins and expose AMR ligands (galactose) *(36, 37)*, and we confirmed its upregulation in response to SA challenge by flow cytometry (**fig. S8B**). Probing the observed therapeutic effect of P2Y12 inhibition in this context, we found ticagrelor strongly inhibited SA-induced platelet sialidase activity (**Fig. 4E**). These results suggest that SA α-toxin-induced thrombocytopenia may not depend on wholescale platelet injury, but instead involve accelerated hepatic AMR-dependent clearance of desialylated platelets upon surface mobilization of Neu1. Binding of ADP to P2Y12 elevates cytosolic calcium (Ca^2+^) levels by stimulating phospholipase C-mediated production of inositol-1,4,5-trisphosphate (IP3), which in turn releases Ca^2+^ from the intracellular stores through IP3 receptor channels. Because Ca^2+^ is a major signaling molecule that allows for ADP-induced lysosomal secretion, P2Y12 inhibition in theory would block this process. However, a singular correlation between Ca^2+^ levels and granular secretion remains ambiguous, as our data, as well as older literature, suggest that there are both Ca^2+^-dependent and Ca^2+^-independent granular secretory pathways *(38, 39)*. That said, by linking sialidase activity to the ticagrelor therapeutic effect, additional target points for pharmacological support of platelet defense against SA came into view.

**Fig. 4.**
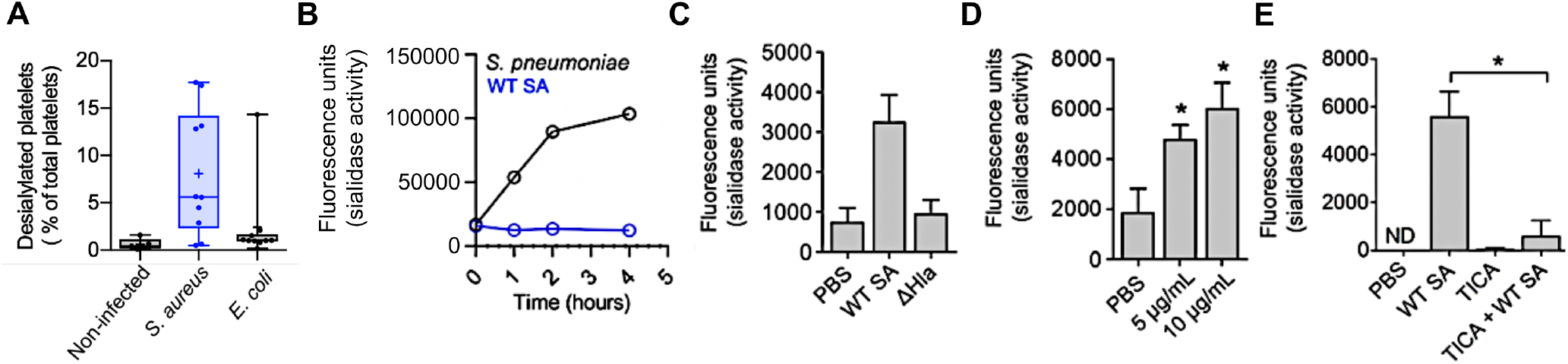
SA α-toxin activates endogenous platelet sialidase activity. **(A)** % desialylated platelets in platelet rich plasma from non-infected control, SA bacteremia or *Escherichia coli* bacteremia patients measured by flow cytometry. **(B)** SA and *S. pneumoniae* sialidase activity assessed for over 4 h. **(C)** Sialidase activity examined on washed human platelets exposed to WT SA or its isogenic ΔHla for 1 h or **(D)** sialidase activity examined on washed human platelets exposed to 5 μg/mL and 10 μg/mL recombinant α-toxin for 30 min. **(E)** Sialidase assay performed on washed human platelets treated with or without 10 μM Ticagrelor and exposed to WT SA for 1 h. Where applicable, all data represented as mean ± SEM and are representative of at least 3 independent experiments. Statistical significance determined by one way ANOVA with Bonferroni’s multiple comparisons test (A, C, D, E). *P < 0.05. PBS, phosphate buffered saline; ns, not significant. ND, not detectable.

### Inhibition of the hepatic Ashwell-Morell receptor (AMR) supports platelet-mediated defense against SA bacteremia

In previous work, we showed that moderate thrombocytopenia mediated by AMR-dependent clearance of desialylated platelets was protective in experimental sepsis caused by *S. pneumoniae*, a sialidase-expressing pathogen *(40, 41)*. However, as shown earlier (**Fig. 1C**), *S. pneumoniae* is resistant to human platelet killing, so removal of desialylated and hypercoagulable platelets does not deplete the bloodstream of an effective antimicrobial effector cell type. We asked whether the innate immune calculus could prove different for platelet-<u>sensitive</u> SA by challenging WT and AMR-deficient (Asgr2^−/−^) mice in the C57bl/6 background. Whereas WT mice remained highly sensitive to SA α-toxin-induced thrombocytopenia, platelet counts in Asgr2^−/−^ mice did not drop following the bacterial challenge (**Fig. 5A**), indicating that recruitment of AMR clearance was the main pathogenic driver of platelet clearance during infection. And in striking contrast to the findings in *S. pneumoniae* infection *(40)*, Asgr2^−/−^ mice, which are resistant to pathogen-induced thrombocytopenia, exhibited a strong survival advantage against lethal SA challenge (**Fig. 5B**). This genetic association could be reproduced pharmacologically in WT mice, where asialofetuin, a competitive glycoprotein inhibitor of the hepatic AMR (**Fig. 5C**), improved mouse survival in lethal SA challenge by maintaining platelet count during infection (**Fig. 5D**) and by reducing bacterial burden in the kidney, livers, and spleen (**Fig. 5E**). Corroborating that α-toxin-dependent desialylation is driving the accelerated platelet clearance, no SA-induced reduction in platelet count was seen in mice lacking the hepatic AMR (**Fig. 5A, Fig. 5F**).

**Fig. 5.**
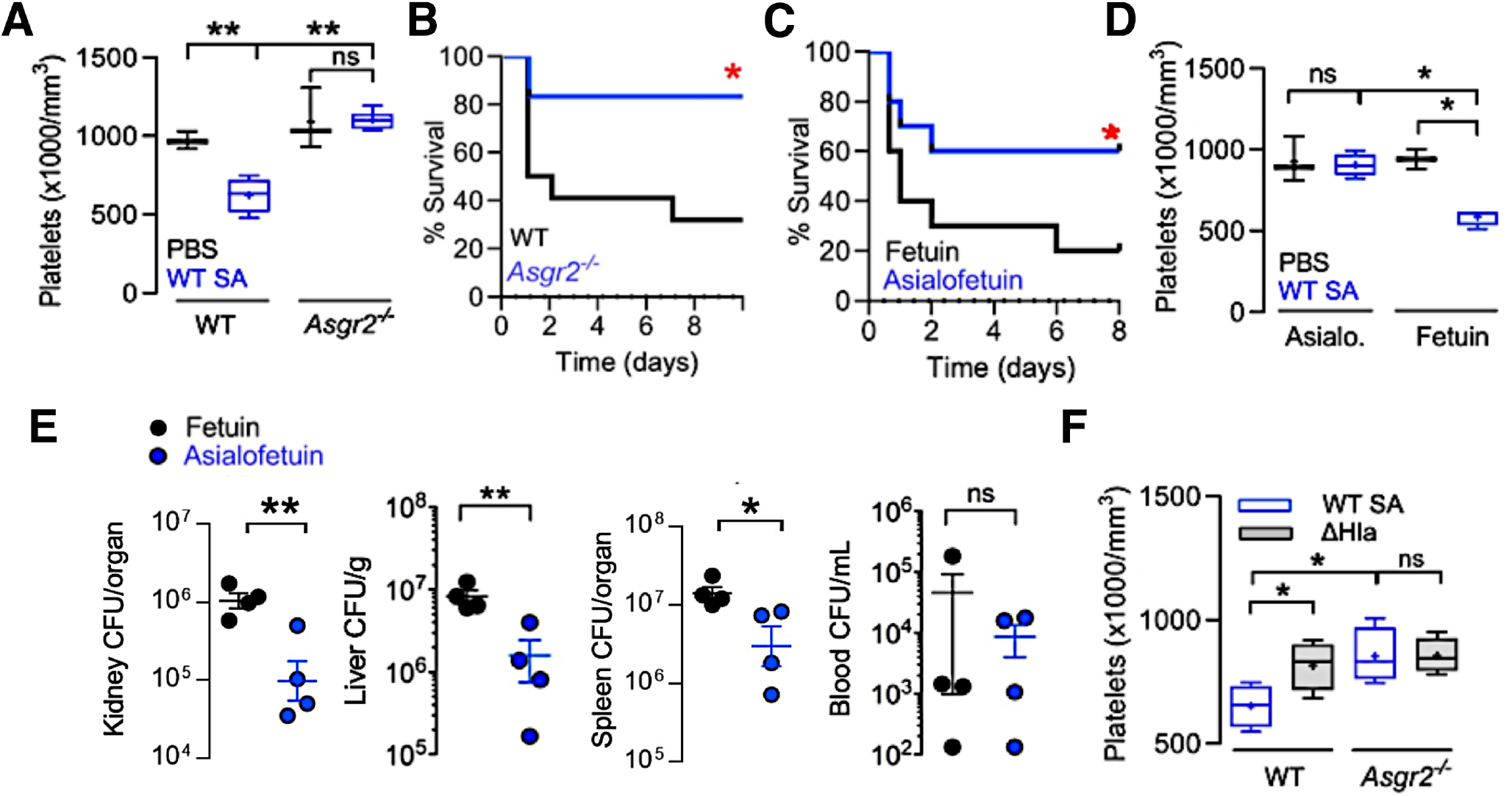
Inhibition of the hepatic Ashwell-Morell receptor (AMR) supports platelet-mediated defense against SA bacteremia. **(A)** C57/Bl6 (n = 4) and Asgr2^−/−^ (n = 6) mice challenged by intraperitoneal injection with SA, blood harvested by cardiac puncture, and platelet count enumerated. **(B)** 10-day mortality study with C57/Bl6 (n = 22) and Asgr2^−/−^ mice (n = 16) challenged by intraperitoneal injection with SA. Study performed two independent times and data pooled. **(C)** 8-day mortality study with C57/Bl6 treated with fetuin (n = 10) or asialofetuin (n = 10) and challenged by intraperitoneal injection with SA. **(D)** C57/Bl6 mice treated with asialofetuin (n = 4) or fetuin (n = 4) and challenged by intraperitoneal injection with SA, platelet count enumerated, and **(E)** kidneys, liver, spleen, and blood harvested 24 h post infection for bacterial colony forming unit enumeration. **(F)** C57/Bl6 and Asgr2^−/−^ mice challenged with wild-type MRSA or the isogenic ΔHla mutant. 4 h post-infection, blood harvested by cardiac puncture for enumeration of platelet count. Statistical significance was determined by unpaired two-tailed Student’s T-test (E), two-way ANOVA with Bonferroni’s multiple comparisons posttest (A,D,F) and Log-rank (Mantel-Cox) Test (B,C) for the survival curves. For floating bar graphs, + denotes the mean, whiskers represent min. to max, and floating box represents 25th to 75th percentile. *P < 0.05, **P < 0.005. PBS, phosphate buffered saline; ns, not significant.

### FDA-approved sialidase inhibitor oseltamivir blocks AMR-mediated platelet clearance and protects against SA bacteremia

The above results showed that mice were protected against SA infection by ticagrelor, which inhibits α-toxin-induced platelet desialylation, or by genetic or pharmacological inactivation of the AMR, which blocks hepatic clearance of desialylated platelets. Further corroboration of the importance of platelet sialylation for maintaining bloodstream defense against SA bacteremia was obtained using mice lacking the St3gal4 sialyltransferase gene, which show diminished platelet sialylation and baseline thrombocytopenia *(42)*. Compared to WT C57bl6 mice, the St3gal4^−/−^ mice had accelerated mortality upon IV SA infection (**Fig. 6A**), but no further SA-induced reduction in their already low platelet counts (~25% of normal mice, **Fig. 6B**). Since sialidase (Neu1) activity appears central to the “toxin-platelet-AMR” pathway driving deleterious thrombocytopenia in SA bloodstream infection, we considered the possibility that pharmacological sialidase inhibition could be of therapeutic benefit. Oseltamivir (Tamiflu^®^) is a commonly prescribed FDA-approved drug designed to target influenza sialidase (neuraminidase) and lessen the severity of flu symptoms. However, oseltamivir has a degree of non-selectivity in its sialidase inhibition, as the drug was recently recognized to raise platelet counts in mice with anti-GPIbα-mediated thrombocytopenia *(43)*. Using Asgr2^−/−^ mice to prevent immediate clearance of desialylated platelets, we confirmed that oseltamivir inhibits platelet desialylation *in vivo* during SA infection (**Fig. 6C**). Then using WT mice, we showed that oseltamivir significantly reduced the degree of α-toxin-induced thrombocytopenia during WT SA infection (**Fig. 6D**), and both oseltamivir and established human Neu1-selective sialidase inhibitor C9-butyl-amide-2-deoxy-2,3-dehydro-N-acetylneuraminic acid (DANA), significantly improved survival outcomes in lethal SA bacteremia (**Fig. 6E**).

**Fig. 6.**
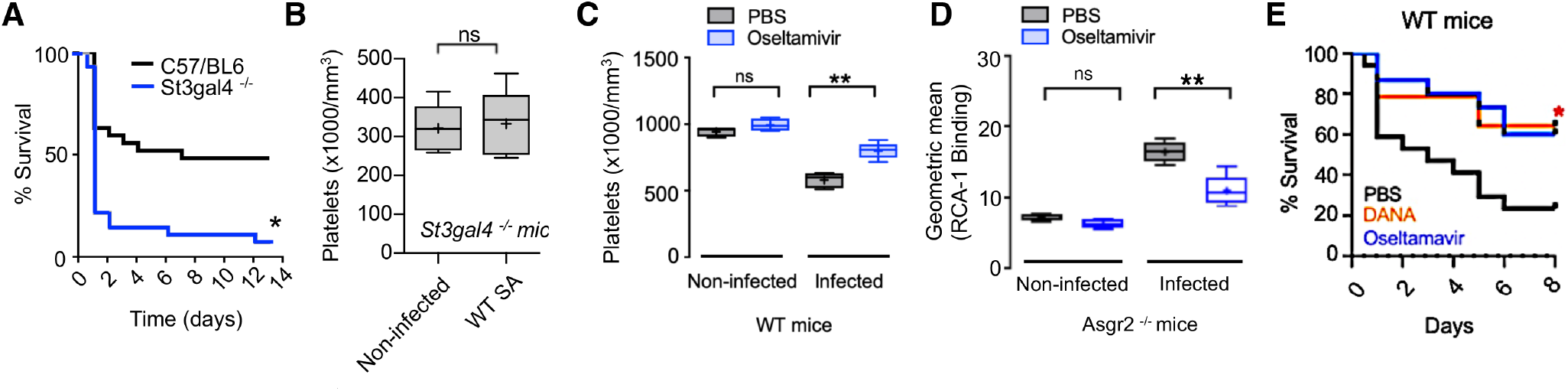
FDA-approved sialidase inhibitor oseltamivir blocks AMR-mediated platelet clearance and protects against SA bacteremia. **(A)** St3gal4^−/−^ mice which have decreased platelet sialylation and thrombocytopenia show accelerated mortality upon SA bloodstream infection (n = 10 per group). (B) No difference in circulating platelet count 4 h after IV SA challenge in WT vs. St3gal4^−/−^ mice (n = 10 per group). **(C)** Platelets isolated from Asgr2^−/−^ mice treated with or without oseltamavir and infected with MRSA were assessed for RCA-1 lectin binding. **(D)** C57/Bl6 mice treated with oseltamavir (n = 6) or PBS control (n = 5) and infected with WT SA by intraperitoneal injection. Blood harvested 24 h after infection and platelet counts collected. **(DE)** 8-day mortality study conducted on C57/Bl6 mice treated with DANA (n = 16), oseltamavir (n = 16), or PBS control (n = 16). Statistical significance was determined by unpaired two-tailed Student’s T-test (B), two-way ANOVA with Bonferroni’s multiple comparisons posttest (C,D) and Log-rank (Mantel-Cox) Test (A,E). Where applicable, statistical significance determined by unpaired Two-tailed Student’s T-test. For floating bar graphs, + denotes the mean, whiskers represent min. to max, and floating box represents 25th to 75th percentile. Unless otherwise stated, *P < 0.05, **P < 0.005. PBS, phosphate buffered saline; ns, not significant.

Finally, to support all elements of the elucidated “toxin-platelet-AMR” pathway, we repeated several key *in vivo* experiments (the ticagrelor, oseltamivir and asialofetuin treatment studies as well as challenge of Asgr2^−/−^ mice) with the SA ΔHla knockout mutant. Given the attenuated virulence of this mutant, establishment of bacteremia in the murine IV model required an 8-fold higher challenge inoculum. Blood harvested 4 h post infection in all the experiments showed that the platelet count drop associated with WT infection was not seen in ΔHla-infected mice in any of the models, even at the 8-fold higher bacterial challenge inoculum (**fig. S9A**). Likewise, equal CFU counts of the ΔHla knockout mutant SA were recovered in the kidney, liver, spleen, and blood of control mice vs. mice treated with either TICA, asialofetuin, or oseltamivir, as well as WT mice when compared to Asgr2^−/−^ animals (**fig. S9B**). As predicted by the model, deletion of α-toxin “phenocopies” the therapeutic benefits of the drug and genetic interventions to support platelet defense against SA bloodstream infection.

## DISCUSSION

SA is a leading agent of human bloodstream infection, with higher morbidity and mortality rates than other common bacterial pathogens. Successful treatment of SA bacteremia remains vexing, as the pathogen deploys diverse mechanisms for resistance to immune and antibiotic clearance. Life-threatening complications of SA bacteremia, such as metastatic infections, infective endocarditis, and disseminated intravascular coagulation, drive worsened patient outcomes. Thrombocytopenia (platelet count <150,000/mm^3^ blood) is a common phenotype observed during bacteremia and is the most predictive independent risk factor for bacteremia-associated mortality, especially in cases of neonatal sepsis and critically ill septic patients in the intensive care unit *(44, 45)*. While the underlying cause of thrombocytopenia is multifactorial, our mechanistic analysis of platelet-mediated defense establishes a central pathophysiologic framework aggravating platelet depletion during SA bacteremia. We show that for SA to evade platelet microbicidal activity, the pathogen deploys the cytotoxic α-toxin, which injures platelets and stimulates the release of endogenous sialidase, thereby dysregulating the platelet clearance mechanism involving the hepatic AMR. Pharmacological targeting of multiple levels of this “toxin-platelet-AMR” pathway reveal new strategies to mitigate the progression of this immunocompromised state and protect against lethal SA bacteremia (**Fig. 7**).

**Fig. 7.**
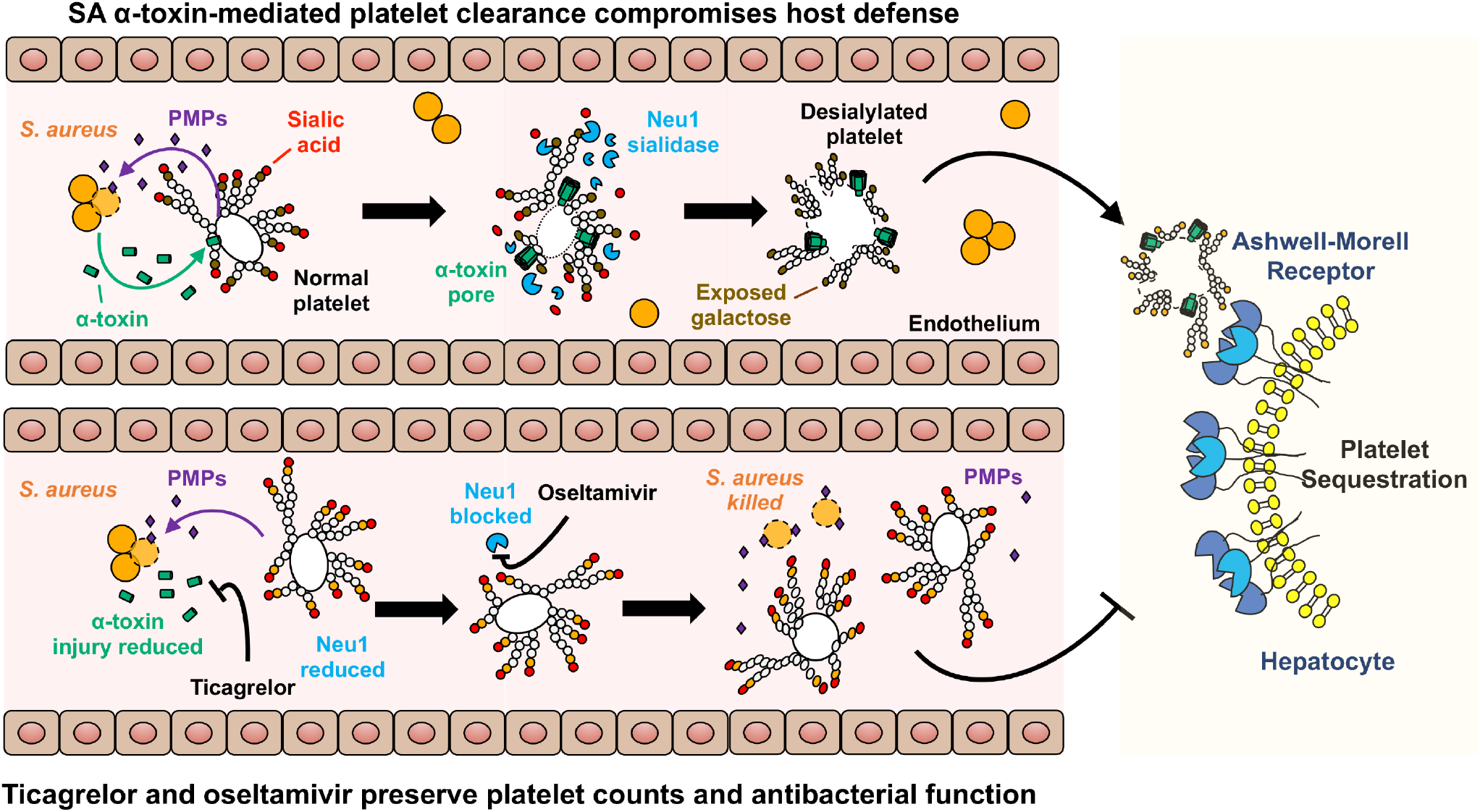
Schematic illustration of proposed “toxin-platelet-AMR” pathway exploited by SA in the pathogenesis of bloodstream infection. Our experiments suggest the possibility of therapeutic drug repurposing of P_2_Y_12_ (e.g. ticagrelor) and sialidase inhibitor (e.g. oseltamivir) drugs to maintain platelet homeostasis and enhance innate immune clearance of the pathogen.

Sialidase transfer from lysosomal compartments to the platelet cell surface may potentially be elicited by multiple surface-bound receptors including but not limited to: P2Y12, PAR-1, and PAR-4 *(46)*. These receptors have converging intracellular signaling pathways, and studies indicate that P2Y12 functions in crosstalk with PAR receptors *(47)*. Ticagrelor, a commonly prescribed FDA-approved P2Y12 inhibitor, hastened clearance of SA bacteremia *in vivo* and enhanced human platelet killing of SA *ex vivo*. Here we describe new effects of the drug in reducing SA α-toxin-mediated platelet cytotoxicity, inhibiting activation of endogenous platelet sialidase activity, and preventing AMR-dependent platelet clearance. As thrombocytopenia was not observed in SA-challenged mice lacking the AMR, wholescale SA-induced platelet damage, and ticagrelor’s ability to counteract this injury, may be a phenomenon unique to the high bacterial concentrations and close platelet contact present in our *in vitro* assays, perhaps only relevant *in vivo* within an infected thrombus. Conceivably, ticagrelor’s primary therapeutic indication for acute coronary syndrome, reduction of platelet aggregation, may further provide protective benefit in SA bacteremia, since the pathogen produces two clotting factors, coagulase (Coa) and von Willebrand factor binding protein (vWbp), that contribute to abscess formation and systemic virulence *(48)*. Of note, one prior report linked *in vitro* P2Y12 activation to the release of platelet antimicrobial peptides active against SA *(49)*. This paradoxical result may in part reflect the particular SA strain (ISP479C) used in the study, which harbors a chromosomal Tn551 insertion with a pleiotropic effect on several extracellular and cell wall proteins, including elimination of measurable α-toxin activity *(50)*. Platelet release of antimicrobial peptides active against SA is also activated by additional pathways, including thrombin-mediated enzymatic activation of cell surface protease-activated receptor-1 (PAR-1) still operative during P2Y12 blockade *(12)*. The feasibility of ticagrelor as an adjunctive therapy for SA bacteremia in complex ICU patients is likely enhanced by its reversible binding to the P2Y12 receptor binding providing a very rapid onset and offset of action *(51)*.

Oseltamivir, a commonly used FDA-approved influenza sialidase inhibitor, maintained platelet sialylation levels to delay AMR-dependent platelet clearance, and thus provided protection against mortality in SA bloodstream infection. While one enzymatic study suggested that oseltamivir had only limited inhibitory activity against human sialidases *(52)*, humans prescribed oseltamivir show higher platelet counts than matched controls (independent of proven influenza, *(53)*), and two independent case studies report successful use of the drug to restore platelet counts in a patient with immune thrombocytopenia *(54, 55)*. Bacterial co-infection is estimated to have contributed to nearly all influenza deaths in the 1918 influenza pandemic and up to one-third of 2009 pandemic influenza A(H1N1) infections managed in ICUs worldwide *(56)*. In particular, the potential for lethal synergism between SA and influenza virus has recently been documented in U.S. clinical epidemiologic studies of adult and pediatric patients *(57, 58)*. In laboratory-confirmed influenza, an inverse relationship between virus load and platelet count is seen, and viral-induced thrombocytopenia can be recapitulated in the ferret model *(59)*. We speculate that the “two-hit” scenario of influenza neuraminidase on top of α-toxin-induced endogenous sialidase activation may accelerate platelet clearance, depleting the host of a critical frontline defense against SA bloodstream dissemination, thus increasing the odds of fatal outcome.

An important limitation of pharmacological targeting AMR-dependent clearance of desialylated platelets to treat bacteremia is its dependency on sensitivity of the offending pathogen to platelet antimicrobial activity. A definitive or strong presumptive microbiologic diagnosis of SA would be required, precluding its use as empiric therapy wherein other pathogens, such as platelet-resistant *S. pneumoniae*, could yield adverse results *(41)*. Multiple pathogenic mechanisms contribute to sepsis and intrinsic host factors can have differing roles depending on the pathogen involved *(60)*. In this case, AMR function may serve protective and disadvantageous roles depending upon the pathogen and the balance of platelet action in thrombosis vs. antimicrobial activity. Prior research indicates that loss of AMR can increase platelet count and regulate thrombopoietin production *(61)*. Mechanisms of physiologic platelet turnover remain to be fully established and are likely to contribute to therapeutic modulation in the future. However, it is unlikely that either ticagrelor or oseltamivir administered late in the course of severe SA-induced thrombocytopenia could quickly restore platelet counts. There, perhaps platelet transfusion could augment anti-SA killing capacity in blood, wherein the pharmacological agents could mitigate against further α-toxin driven accelerated desialylation and AMR clearance of the donor platelets. The pathological process can also be targeted upstream at the level of the inciting SA α-toxin, where important research on neutralizing antibodies (e.g. Medimmune 4893) and receptor antagonists (e.g. GI254023X)) have shown promising results *(14, 62)*.

Therapeutic drug repurposing is an important avenue of exploration to improve clinical outcomes in serious infections where high rates of treatment failure and antibiotic resistance jeopardize patients. Elucidation of sialidase-dependent platelet homeostasis as a key battleground in host defense against SA bloodstream infection revealed the potential utility of P2Y12 and sialidase inhibition as adjunctive agents to antibiotic treatment and ICU supportive care for the critically ill. The most effective physiological concentrations to inhibit platelet cytotoxicity, sialidase activity and protect against SA bacteremia in humans are currently unknown. As FDA-approved drugs with excellent safety profiles in each class are readily at hand, we hope that carefully designed clinical investigation to validate or refute our experimental observations may follow.

## MATERIALS AND METHODS

### Study Design

The objective of this study was to understand the mechanistic basis of platelet homeostasis and function during SA bacteremia to guide future therapeutic approaches. Our analysis of patient data and SA isolates from a published 2009-10 IRB-approved study *(25)* of SA bacteremia at the University of Wisconsin Hospital linked thrombocytopenia to patient mortality and elevated α-toxin production. Both correlations were corroborated in a UC San Diego IACUC-approved murine model of SA bacteremia. UC San Diego IRB-approved *ex vivo* studies with freshly isolated human platelets were used to show the FDA-approved P2Y12 antagonist ticagrelor blocked α-toxin induced platelet injury and sialidase activation, improving microbial killing. Infection studies WT and isogenic AMR-deficient mice were used to link α-toxin mediated platelet sialidase activation to accelerated thrombocytopenia and impaired SA clearance, which could be counteracted by ticagrelor or the FDA-approved sialidase inhibitor oseltamivir.

### Statistical Analysis

All *in vitro*, *ex vivo*, and *in vivo* data were collected from three or more (≥3) independent experiments with ≥3 biological replicates and are represented as mean ± standard error mean (SEM), unless otherwise stated. For descriptive data (transmission electron microscopy and histopathologic staining), experiments were performed at least twice independently with ≥3 biological replicates and illustrated as best representative images. The alpha level used for all tests was 0.05; unpaired Student’s t-test, one-way ANOVA with Bonferroni’s multiple comparisons test or two-way ANOVA with Bonferroni’s multiple comparisons test was performed as explained in figure legends to determine statistical significance. For comparison of survival curves, a log-rank (Mantel-Cox) test was performed. Statistical analyses were done using GraphPad Prism, version 8.42 (GraphPad Software Inc., La Jolla, CA, USA). P values < 0.05 were considered statistically significant.

## ACKNOWLEDGEMENTS

This work was supported by NIH grants HL125352 (JDM, VN), HL107150 (VN), KD048247 (JDM), AI124326 (GS, VN), HD090259 (GS, VN) and AI13262 (WR). JS was supported the UCSD PharmD/PhD Program and IC by the UCSD Research Training Program for Veterinarians (NIH OD017863). The authors thank Ajit Varki for his critical insights on human and murine sialobiology.

## Supplementary Materials

### MATERIALS AND METHODS

#### Ethics statement

Animal studies were conducted in accord with protocols approved by the UC San Diego Institutional Animal Care and Use Committee; all efforts were made to minimize animal numbers and suffering. Blood for platelet isolation was obtained via venipuncture from healthy volunteers under written informed consent approved by the UC San Diego Human Research Protection Program.

#### *S. aureus* patient isolates

Consecutive patients from previously published, IRB-approved study *(63)* and its ongoing continuation (IRB #2018-0098) with blood cultures of *S. aureus* (SA) from April 2009 through March 2010 at the University of Wisconsin Hospital (a 493-bed academic medical center in Madison, WI) were analyzed for α-toxin expression by western immunoblot and densitometry band analysis by Image J. Levels of α-toxin expression was grouped in the following order: low: >10,000; medium: 10,000 – 20,000; high: >20,000. Patient demographics, blood work (including platelet and leukocyte counts), and infection source were collected at time of administration. Bacterial isolates obtained at the onset of presentation and stored at −80°C until analysis. All laboratory tests were performed by investigators blinded to patient information.

#### Measurement of desialylation of patient platelets

In the above patient biobank, 2-5 mL of patient whole blood is collected on the day of presentation with bacteremia at the hospital. In this analysis, platelets from patients with SA bacteremia were compared to patients with *Escherichia coli* bacteremia and sepsis, and uninfected healthy volunteers. All patients with *S. aureus* were diagnosed with bacteremia from an endovascular source by an infectious diseases physician. Patients with *E. coli* bacteremia/sepsis had infection sources from the urinary or gastrointestinal tract. No patients received P2Y12 antagonists or any other anti-platelet therapy at the time of collection. Samples were processed to separate plasma and cells, which are then aliquoted separately stored at −80°C until analysis. Platelets in plasma from patients admitted with SA, *Escherichia coli* sepsis, or normal controls were analyzed for RCA-1 binding by flow cytometry. Plasma samples were stained with PerCP/Cy5.5 anti-human CD41 antibody (Biolegend) and Fluorescein labeled Ricinus Communis Agglutinin I (RCA-1 : Vector Laboratories). % desialylated platelets were calculated using the Flojo software.

#### Bacterial strains and plasmids

Community-acquired methicillin-resistant SA (MRSA) strain USA300 (TCH1516) and its isogenic ΔHla mutant lacking α-toxin, were used in the study. Targeted mutagenesis of *Hla* was conducted by precise, markerless allelic replacement of USA300 TCH1516 *hla* gene (Locus tag USA300HOU_1099, NC_010079.1 (1170314.1171273, complement) by PCR-based methods adapting the pKOR1 knock-out strategy previously described for SA mutagenesis *(64)*. As shown in fig. S3, sequence immediately upstream of *hla* was amplified with the primers D and E and that immediately downstream of *hla* with the primers B and F. Primers B and D were constructed with ~25 bp 5’ overhangs for the opposite flanking region. Upstream and downstream PCR products were fused using primers E and F in a second round of PCR. The amplicons were then subcloned into temperature-sensitive vector pKOR1 using the BP clonase reaction (Invitrogen). The resulting plasmid pKOR1-hla was passed through SA RN4220, and the purified plasmid electroporated into strain TCH1516. Precise in-frame allelic replacement of *hla* was established by a two-step process of temperature shifting and anti-sense counterselection and confirmed by PCR. Other listed primers (**fig. S3**) were used for PCR confirmations and for screening potential mutants and clones. All strains were routinely grown in Todd Hewitt broth (THB) and propagated shaking at 37°C to mid-log phase (optical density 600 nm (OD_600_) = 0.4) unless otherwise stated. Bacteria were collected by centrifugation at 4,000 RPM x 10 min, washed once in phosphate-buffered saline (PBS), and resuspended to the desired dilution in PBS.

#### Platelet and neutrophil isolation

For platelet isolation, human venous blood was drawn using a 20G needle from healthy human donors using acid-citrate-dextrose buffer (ACD; Sigma) as an anticoagulant (1:6 v/v), unless otherwise stated. To obtain platelet-rich plasma (PRP), blood was centrifuged at 1,000 RPM x 10 min with no brake. To avoid contaminations with other cell types, only the upper two thirds of the platelet-rich plasma fractions were used. PRP was centrifuged at 1.500 RPM x 10 min. Isolated platelets were resuspended in serum-free, antibiotic-free, inhibitor-free RPMI (without phenol red) at room temperature. Blood was drawn according to a protocol approved by the local ethics committee. For neutrophil isolation, venous blood was drawn from healthy donors as above but using heparin as an anticoagulant. Purified neutrophils were collected using Polymorph Prep (Axis-Shield, Dundee, Scotland) per manufacturer’s protocol.

#### Platelet cytotoxicity

Human platelets were pre-treated with 10 μM ticagrelor (Sigma) or vehicle control and incubated with rotation at 37° C + 5% CO_2_ for 20 min, then exposed to 5 μg/mL recombinant α-toxin (H9395 Sigma) for an additional 30 min. Samples were spun down at 500 x *g* for 5 min, and supernatants evaluated using a commercial lactate dehydrogenase (LDH) assay (Promega) or ATP assay (Promega, CellTiter-Glo® Luminescent Cell Viability Assay).

#### Human platelet sialidase (neuraminidase) assay

Washed human platelets (3 × 10^7^) were assayed for sialidase activity using a slight modification of the previously published 2′-(4-methylumbelliferyl)-α-D-*N*-acetylneuraminic acid (4MU; Sigma) assay *(65, 66)*. Platelets were added to a white 96-well plate (Costar), pretreated with or without the identified drugs for 15 min, and exposed to 3 × 10^7^ CFU of SA (MOI = 1:1) for 1 h at 37°C in 5% CO_2_. Next, 125 μM 4MU was added and incubated 30 min at 37°C before 1M Na_2_CO_3_ was added to each sample and fluorescence determined at excitation 530 nm and emission 585 nm. Background fluorescence was measured following the same procedure but in the presence of 1 mM sialidase inhibitor DANA.

#### Human platelet ADAM10 protease assay

Human platelets were isolated from healthy donors using hirudin as an anticoagulant. Isolated platelets were resuspended in serum-free, antibiotic-free, inhibitor-free RPMI (without phenol red) at room temperature. Human platelets (2 × 10^7^) were pre-treated with 10μM ticagrelor (Sigma) or vehicle control and incubated at 37°C with rotation for 20 min, at which time platelets were exposed to 5 mg/mL recombinant α-toxin and a fluorogenic ADAM10 specific substrate (PEPMCA001, Biozyme) at 37°C + 5% CO2 without shaking. Fluorescence was measured every 15 min for 2.5 h using wavelengths of 325 nm (excitation) and 393 nm (emission).

#### Calcium assay (Fluo-3 assay)

Human donor blood was collected using hirudin as an anticoagulant. Platelet-rich plasma (PRP) was isolated and incubated with 2mM Fluo-3 AM (Thermo Fisher Scientific) at 37°C + 5% CO2 with rotation for 20 min. Isolated platelets were resuspended in serum-free, antibiotic-free, inhibitor-free RPMI (without phenol red) at room temperature, pre-treated with 10μM Ticagrelor (Sigma) or vehicle control for 15 min, then exposed to 5 μg/mL recombinant α-toxin. Fluorescence was measured at 505 nm excitation and 530 nm emission every 30 sec.

#### Transmission electron microscopy

Isolated human platelets were pre-treated with 10 μM ticagrelor (Sigma) or vehicle control at 37°C + 5% CO2 for 20 min, then infected with MRSA at MOI = 0.01 for an additional 2 h. Samples were fixed in modified Karnovsky’s fixative (2.5% glutaraldehyde + 2% paraformaldehyde in 0.15 M sodium cacodylate buffer, pH 7.4) for at least 4 h, post-fixed in 1% osmium tetroxide in 0.15 M cacodylate buffer for 1 h, and stained in block in 2% uranyl acetate for 1 h. Samples were dehydrated in ethanol, embedded in Durcupan epoxy resin (Sigma-Aldrich), sectioned at 50–60 nm on a Leica UCT ultramicrotome, and picked up on Formvar and carbon-coated copper grids. Sections were stained with 2% uranyl acetate for 5 min and Sato’s lead stain for 1 min. Grids were viewed using a Tecnai G2 Spirit BioTWIN transmission electron microscope and photographs were taken with an Eagle 4k HS digital camera (FEI). Images were taken from multiple random fields at 1200 ×, 2900 ×, 23,000 ×; gross morphology was analyzed in a blinded fashion.

#### Bacterial growth curves

Sterile non-pyrogenic tubes containing THB treated with ticagrelor (10 μM) or untreated were inoculated with overnight bacterial cultures to achieve an optical density (600 nm) of 0.1. Tubes were incubated in a shaking 37°C incubator, and absorbance measured every 30 min (600nm) for 8 h using a Spectronic 20D+ spectrophotometer (Thermo Scientific, Waltham, MA, USA)

#### Bactericidal assays

*Human platelet killing*: Isolated platelets were pre-treated with 10μM ticagrelor (Sigma) or vehicle control for 20 min at 37°C + 5% CO for 20 min, then infected with MRSA at MOI = 0.01 for 2 h. Infected platelets were sonicated (Fisher Sonic Dismembrator 550) for 3 sec, serially diluted, and plated THA plates. Percent killing by platelets as surviving CFU vs. original inoculum. *Human neutrophil killing:* Freshly isolated human neutrophils (PolymorphPrep) in serum-free RPMI were added to 96-well plates at 5 × 10^4^ cells per well and treated with ticagrelor (10 μM) or vehicle control for 1 h at 37 °C + 5% CO_2_, were infected with MRSA at MOI = 0.01 for 2 h. After incubation, cells were lysed with 0.025% Triton X-100, serial diluted, and plated on THA for CFU enumeration and determination of percent killing vs. inoculum. *THP-1 macrophage killing:* The THP-1 monocyte cell line authenticated by ATCC were cultured in RPMI (with phenol red) medium + 10% fetal bovine serum (FBS). Cells were differentiated in a 96 well format for 48 h with 25 nM phorbol myristate acetate (PMA, Sigma) with a subsequent 24 h rest period in RPMI + 10% FBS. On the day of infection, cells were washed once with PBS, treated with ticagrelor (10 μM) or untreated control for 1 h, and infected with MRSA at MOI 1:100. After incubation, cells were lysed with 0.025% triton X-100, serial diluted, and plated on THA for CFU enumeration and determination of percent killing vs. inoculum. *Human whole blood killing:* Whole blood was drawn from healthy donors using anticoagulant citrate dextrose solution (ACD, 1:6). Blood was pre-treated with varying concentrations of ticagrelor or vehicle control at 37°C + 5% CO_2_ with rotation, then infected with MRSA (1:10) and incubated for an additional 1 h. After incubation, samples were sonicated (Fisher Sonic Dismembrator 550) twice at 20% maximum power for 3 sec with 10 seconds interval, serially diluted, and plated on THA for CFU enumeration and determination of percent killing vs. inoculum.

#### Induction and quantification of neutrophil extracellular traps

To induce extracellular trap production, neutrophils were seeded in 96-well plates at a density of 5 × 10^4^ cells per well in RPMI (without phenol red). Cells were incubated with ticagrelor (10 uM) or untreated control at 37°C with 5% CO_2_ for 1 h before addition of NET-inducing PMA (25nM) for an additional. Extracellular DNA content was quantified using a Quant-IT PicoGreen dsDNA Assay Iit (Life Technologies, Carlsbad, CA) per manufacturer’s instructions.

#### Human platelet surface P-selectin, ADAM-10, GP6 and CD63 measurements

The surface expression of P-selectin, ADAM-10, GP6 and CD63 were measured by flow cytometry. Human platelets were first incubated with TICA or vehicle control for 20 min in 37°C. After a wash with PBS, 1 × 10^7^ platelets were incubated with SA, α-toxin or control media for 90 min. In each tube, antibodies against P-selectin, ADAM-10, GP6 or CD63 each conjugated with PE (Biolegend, San Diego) were added. Expression of each molecule on the platelet surface was detected using FACSCalibur (BD) and analyzed using Flo Jo software (Flojo LLC).

#### Human platelet β-galactosidase activity

Human platelets were incubated with RPMI only, SA or SA and ticagrelor. After 90 min incubation at 37°C, the samples were centrifuged at 2000 rpm for 10 min to collect supernatants. β-galactosidase activity in platelet supernatants was measured using the Mammalian β-Galactosidase Assay (Thermo-Fisher) per the manufacturer’s protocol.

#### Mouse infection and platelet count determination

Wild-type SA and isogenic ΔHla mutant cultures were grown shaking overnight at 37°C in THB, washed once in 1x PBS, and 1 × 10^8^ colony forming units (CFU) were injected intravenously (i.v.) into outbred 8- to 10-week-old CD1 mice (Charles Rivers Laboratories). Blood was collected, platelet count determined, and samples serial diluted and plated onto Todd Hewitt agar (THA) plates for CFU enumeration. Platelet count and CFU burden were enumerated 4 h post infection. For platelet depletion studies, endotoxin and azide-free 1 mg/kg anti-CD41 antibody (clone MWReg30, Biolegend) or 1 mg/kg isotype control rat IgG1 (clone RTK2071, Biolegend) were injected intraperitoneally (i.p.). 1 × 10^8^ SA was administered by tail vein injection (i.v.) 16 h post-antibody treatment, and 4 h later mice were euthanized by CO2 inhalation, blood collected by cardiac puncture with a 25G needle attached to a syringe containing 100 mL ACD buffer, and a complete blood count (CBC) was obtained. Blood, liver, spleen and kidneys were harvested, homogenized, and plated in serial dilutions onto THA plates for CFU counts. For *ex vivo* platelet depletion analysis, mice were treated as above and blood collected by cardiac puncture 16 h after antibody injection, and 1 × 10^6^ SA added to the platelet-depleted and control blood and incubated at 37°C with rotation for 1 h prior to dilution plating for CFU counts. For platelet count determination in a time course after infection, 5 × 10^7^ SA were administered i.v. an 50 μl blood collected by submandibular bleeding into EDTA tubes at each time point. Platelet counts were analysed using ProCyte Dx Hematology Analyzer (IDEXX).

#### Measurement of GP6 expression on murine platelets

GP6 expression on murine platelets during SA systemic infection was measured by flow cytometry. Blood collected above for time course platelet count was incubated with anti-mouse GP6 antibody (Emfret). Expression of GP6 and percentage GP6 negative platelets were analyzed using Flo Jo software (Flojo LLC).

#### Analysis of bone marrow megakaryocyte ploidy in murine SA systemic infection

Measurement of megakaryocyte counts in bone marrow and evaluation of megakaryocyte ploidy was performed as described previously *(67)*. In short, BM cells were harvested from femurs of mice 72 h after SA (5 × 10^7^ CFU) i.v. infection (n = 3 in each group). After gently suspending the BM cells in 0.5% BSA PBS with 2 mM EDTA, cells were fixed in 2% paraformaldehyde at 4°C for 2 h, then washed and resuspended in PBS/EDTA with Fc Block for 10 min. Next 1 μg/mL Brilliant Violet 421™ anti-mouse CD41 antibody (Biolegend,), 75 μg/mL propidium iodide (Sigma-Aldrich) and 45 μg/mL RNAse A (Qiagen) were added and incubated for 30 min. Equal volumes of each sample (100 μl) were analyzed by flow cytometry using a NovoCyte (ACEA Biosciences, San Diego, CA). Megakaryocytes were identified as CD41+ cells and megakaryocyte number/ploidy was analyzed for 4 min.

#### Platelet microparticle analysis by flow cytometry

Isolated human platelets were infected with SA strains for various time points. After each time point, samples were collected, centrifuged and the supernatant analyzed using flow cytometry for microparticle content by adding FITC-conjugated anti-CD41 antibody (Biolegend). FITC positive particles smaller than 1 μm latex beads were selected. The absolute count was calculated from the 15 μm beads added to the same samples. For counting platelet microparticles in mouse serum, animals were infected with 1 × 10^8^ SA i.v., blood collected 4 h after infection, and the serum separated. After adding FITC conjugated anti-CD41 antibody, microparticles in serum were analyzed by FACS as described above.

#### *Asgr2^−/−^*, *St3gal4^−/−^* and AMR inhibitor mouse infection studies

Eight- to 12-week-old *Asgr2*^−/−^ mice *(68)* or 10 to 14-week-old *St3gal4*^−/−^ mice on a C57/Bl6 (Jackson Laboratories) genetic background *(69)* and WT mice bred and raised in the same room were used. WT SA was grown overnight shaking at 37°C in THB, washed once in 1x PBS, and 1 × 10^8^ colony forming units (CFU) injected intraperitoneally (i.p.) unless otherwise specified in the Figure Legend, and mortality observed over the course of 10 days. For AMR inhibitor studies, C57/Bl6 mice were treated with 25mg/mL asialofetuin or fetuin prior to i.p. challenge with 1 × 10^8^ CFU SA; mortality was observed over the course of 8 days. For both studies, mice that appeared moribund were euthanized by CO_2_ asphyxiation. Platelet count enumerations were performed 4 h post infection. For AMR inhibitor CFU enumeration, mice were euthanized 24 h post-infection, organs harvested, and dilution plated onto THA.

#### Sialidase inhibitor mouse infection studies

Eight- to 10-week-old wild-type C57/Bl6 mice were treated with oseltamivir (5 mg/kg) in 100 μL PBS or Neu1-selective inhibitor C9-butyl-amide-2-deoxy-2,3-dehydro-N-acetylneuraminic acid (DANA) (2 mg/kg) at the time of- and 3 hours-post intraperitoneal infection with 1 × 10^8^ CFU SA. Platelets were enumerated and sialidase activity assessed using 2′-(4-methylumbelliferyl)-α-D-*N*-acetylneuraminic acid (4MU; Sigma) from blood collected 4 h post infection per using a previously described protocol *(4)*. For *ex vivo* sialidase analysis, murine platelet rich plasma (PRP) was isolated by cardiac puncture with a 25G needle attached to a syringe containing 100 μL ACD and centrifuged at 100 × *g* for 10 min without braking. Following isolation, 25 μL of PRP was added to wells of white 96-well plate (Costar) with 25 μL RPM I+ 125 μM 4MU. The plate was incubated at 37°C + 5% CO_2_ for 30 min, followed by an addition of 1M Na_2_CO_3_, and fluorescence determined at excitation 530 nm and emission 585nm.

#### Ticagrelor treatment mouse infection studies

SA was grown shaking overnight at 37°C in THB, washed once in PBS, and 1 × 10^8^ colony forming units (CFU) injected intravenously (i.v.) into outbred 8- to 10-week-old CD1 (Charles River Laboratories, Wilmington, MA, USA) mice. Where indicated, ticagrelor (4 mg/kg) or vehicle (water) was delivered by oral gavage 24 h prior- and every 24 h post-infection over a course of 10 days. Mice that appeared moribund were euthanized by CO_2_ asphyxiation. For quantification of CFU burden and histological preparation, mice treated with ticagrelor (4 mg/kg) or vehicle (water) 12 h prior- and every 24 h post-intravenous injection of 1 × 10^8^ CFU SA. At 12 h post-infection, two mice from each group were euthanized by CO_2_ asphyxiation and the kidneys, spleen, heart, and liver harvested and fixed 10% neutral-buffered formalin for 24 h, then routinely processed and paraffin-embedded for histological analysis. Five-micron thick hematoxylin and eosin-stained sections of each tissue were examined by a veterinary pathologist blinded to the treatment group. Distinct bacterial colonies visible at 4X magnification were counted in three longitudinal sections of heart and six longitudinal sections of kidney. Lesions related to bacterial infection were described and graded (minimal-1, mild-2, moderate-3, or severe-4) based on degree of tissue damage. At 72 h post infection, remaining surviving mice were euthanized by CO_2_ asphyxiation, blood collected by cardiac puncture, and organs excised. Blood and organ homogenate (MagNA Lyser instrument (Roche Diagnostics Corporation, Indianapolis, IN) were serially diluted in molecular grade H_2_O and plated onto THA for bacterial CFU enumeration. The study was performed three independent times and data from a representative experiment shown. For platelet quantification, mice were treated with ticagrelor (4 mg/kg) or vehicle (water) every 12 h for 72 h prior to i.v. injection of 1 × 10^8^ CFU SA. Four h post-infection, blood was collected by cardiac puncture with a 25G needle attached to a syringe containing 100 mL ACD buffer, transferred into EDTA tubes and a complete blood count (CBC) was obtained.

#### Mouse infection studies with the SA Δ*hla* mutant

Mice treated with ticagrelor, oseltamivir, asialofetuin or Asgr2−/− mice were infected with 8 × 10^8^ SA ΔHla mutant strain i.v. and platelet counts and bacterial burden analyzed as described above.

#### Statistics

All *in vitro*, *ex vivo*, and *in vivo* data were collected from three or more (≥3) independent experiments with ≥3 biological replicates and are represented as mean ± standard error mean (SEM), unless otherwise stated. For descriptive data (histopathologic staining), experiments were performed at least twice independently with ≥3 biological replicates and illustrated as best representative images. The alpha level used for all tests was 0.05; unpaired Student’s t-test, one-way ANOVA with Bonferroni’s multiple comparisons test or two-way ANOVA with Bonferroni’s multiple comparisons test was performed as explained in figure legends to determine statistical significance. Statistical analyses were done using GraphPad Prism, version 8.42 (GraphPad Software Inc., La Jolla, CA, USA). P values < 0.05 were considered statistically significant.

---

**Fig. S1.**
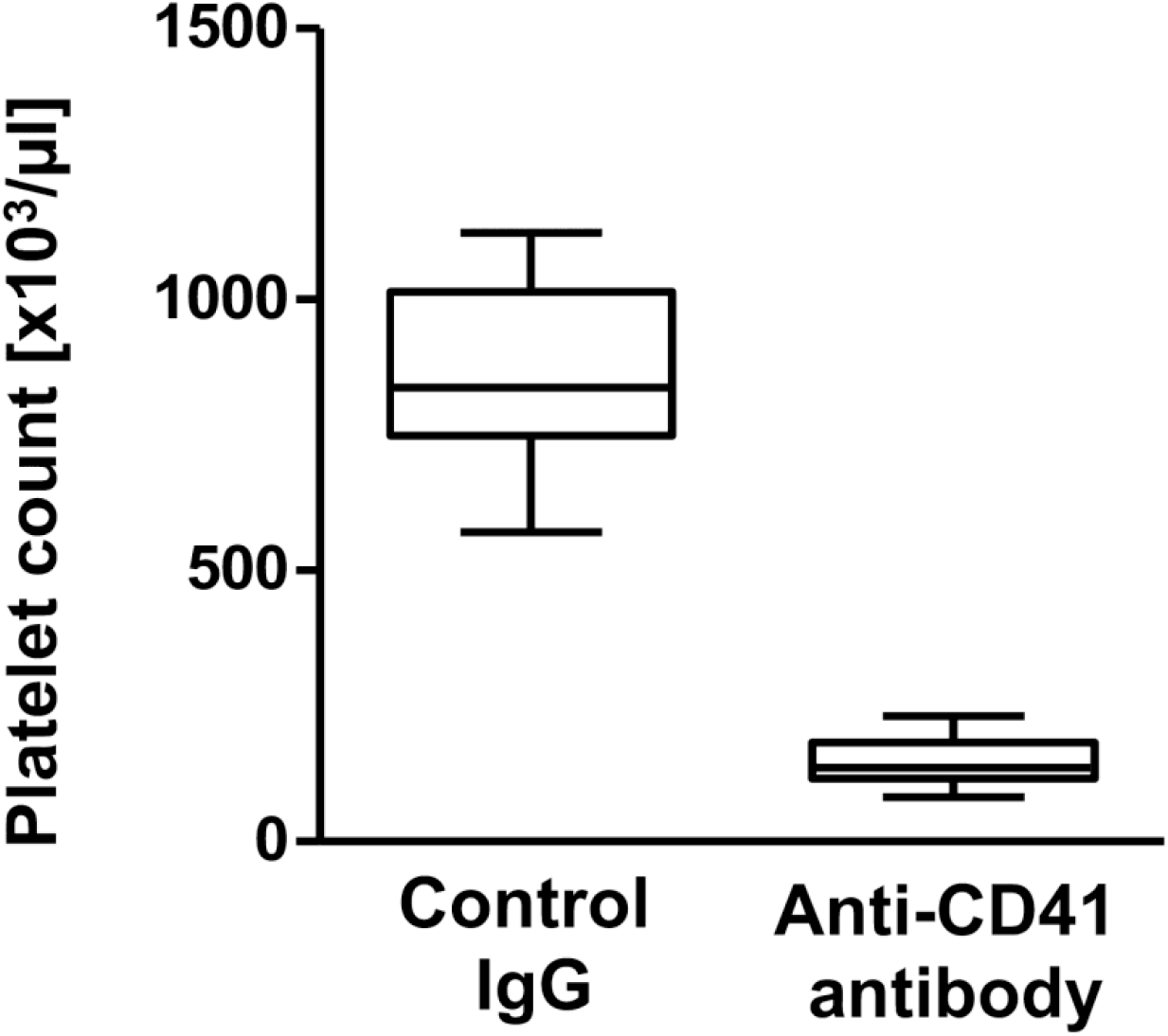
Thrombocytopenia established by anti-CD41 antibody treatment of mice. Endotoxin and azide-free 1 mg/kg anti-CD41 antibody (clone MWReg30, Biolegend) or 1 mg/kg isotype control rat IgG1 (clone RTK2071, Biolegend) were injected intraperitoneally (i.p.) and blood collected by for platelet enumeration 16 h post-antibody treatment.

**Fig. S2.**
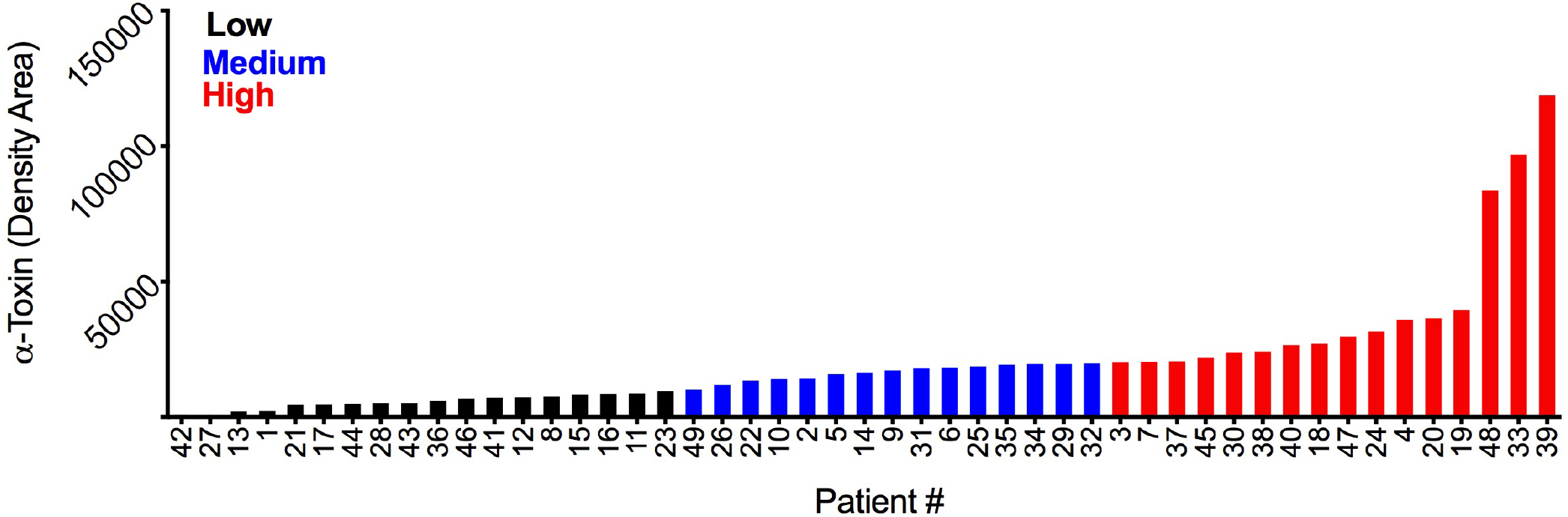
Grouping of α-toxin expression of SA isolates from bacteremia patients. Western immunoblot band analysis on 49 patient clinical isolates grouped according to density area. Groups: low <10,000 (*black*); med. <20,000 (*blue*); high >20,000 (*red*). Densitometry performed using ImageJ software. Data represented as mean only.

**Fig. S3.**
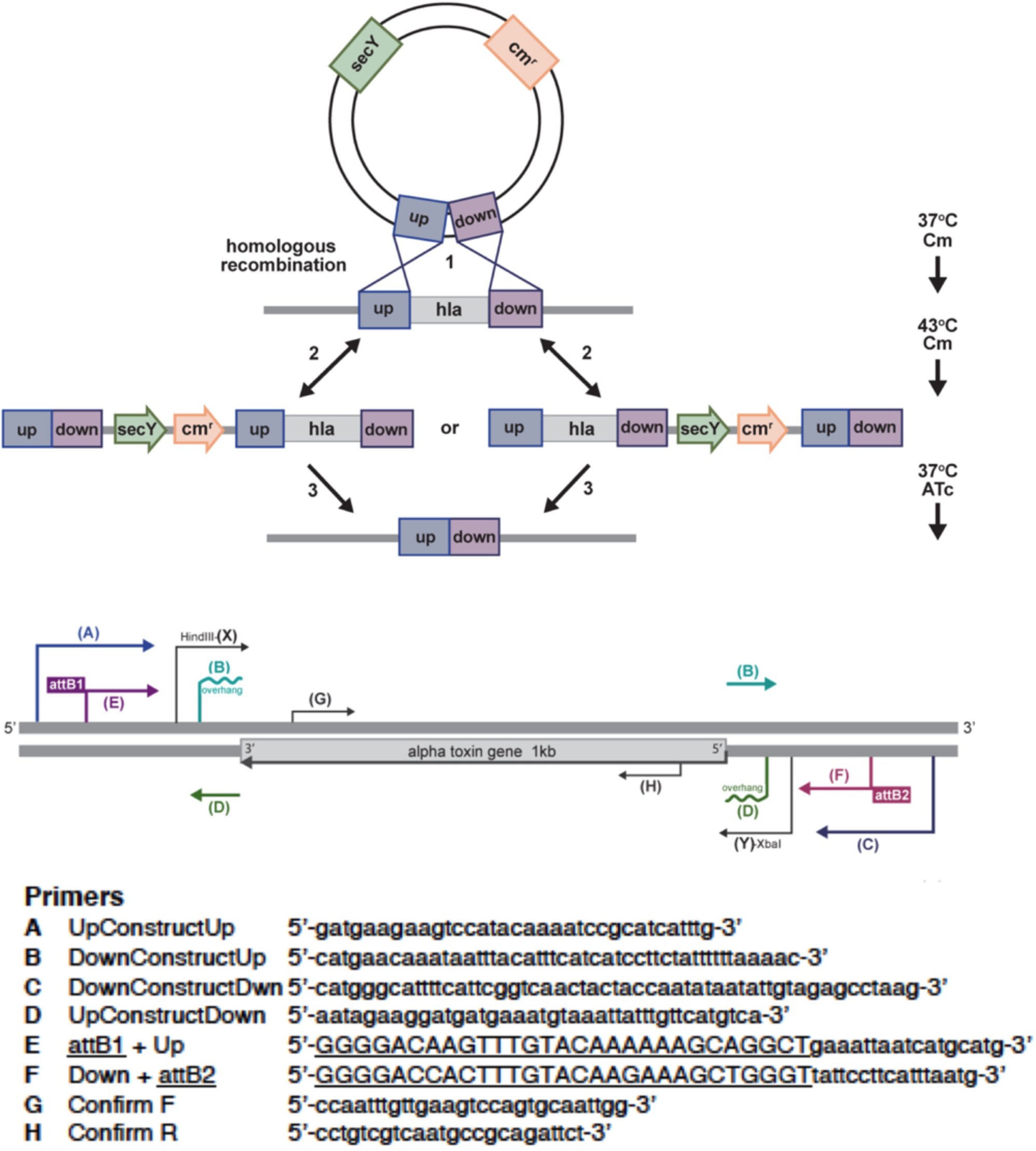
Generation of an isogenic SA ΔHla mutant. Scheme and primer selection for PCR-based targeted markerless allelic replacement mutagenesis of the *hla* gene encoding α-toxin in methicillin-resistant SA strain USA300 TCH1516.

**Fig. S4.**
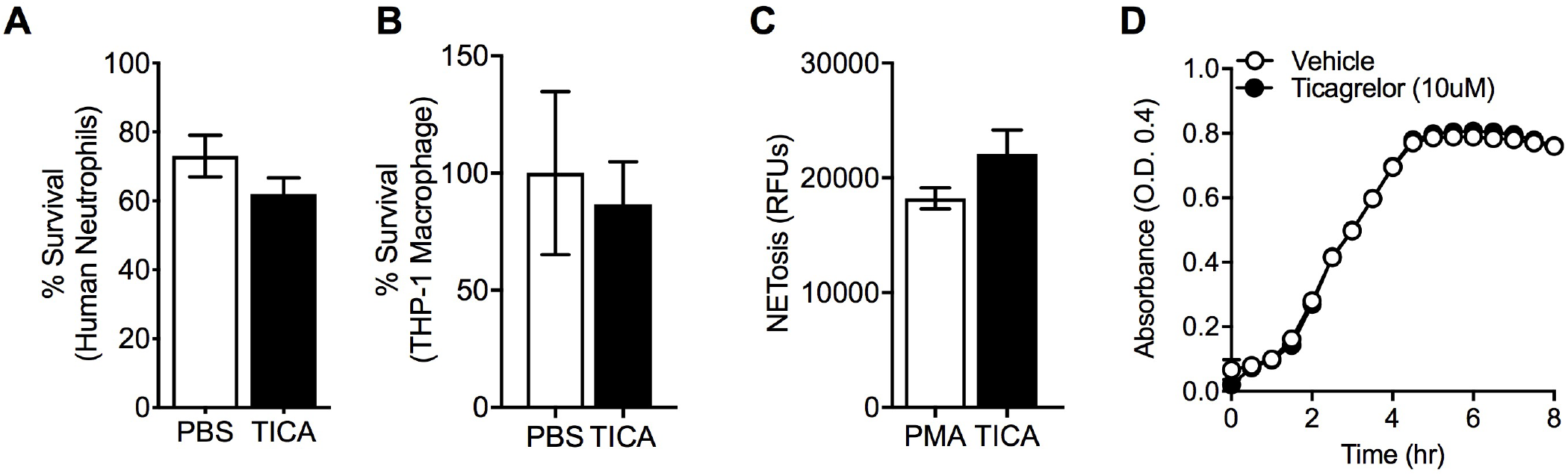
Exclusion of ticagrelor off-target effects on SA immune cell interactions and growth. (**A**) Quantification of MRSA colony-forming units (CFUs) in a purified human neutrophil killing assay. Human neutrophils and (**B**) THP-1-derived macrophages treated with vehicle control (PBS 1x) or 10 μM Ticagrelor for 30 minutes prior to 1 h exposure to MOI: 1 MRSA. (**C**) Isolated human neutrophils pre-treated with vehicle or 10 μM Ticagrelor and subsequently exposed to PMA for quantification of neutrophil extracellular trap (NET) production. (**D**) Growth curve analysis of pre-treated SA with 10μM TICA. Absorbance measured at an optical density (O.D.) of 0.4 measured over the course of 8 hours. Where applicable, data represented as mean ± SEM and are representative of at least three independent experiments. Statistical significance determined by unpaired Two-tailed Student’s t test. \**P* < 0.05

**Fig. S5.**
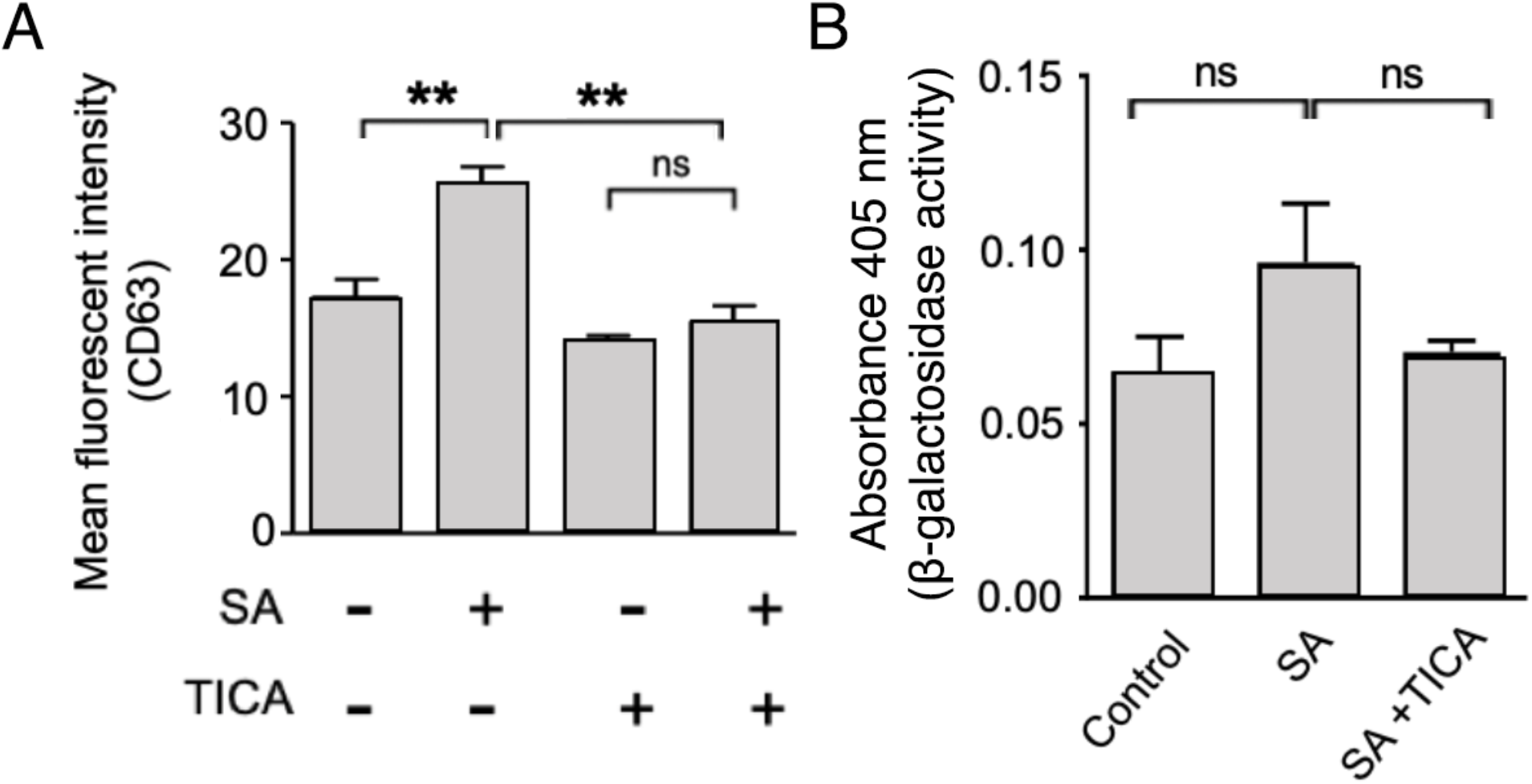
Additional effects of SA or TICA on human platelet phenotypes *in vitro*. (A) Human platelet CD63 expression was measured by flow cytometry with or without TICA treatment and MRSA challenge (MOI = 0.1) after 90 min. (B). β-Galactosidase activity released from human platelets were measured using the colorimetric method of the Mammalian β-Galactosidase Assay Kit (Thermo-Fisher). All data are represented as mean ± SEM and are representative of at least 3 independent experiments. Statistical significance was determined by one-way ANOVA with Bonferroni’s multiple comparisons test (A,C,G), unpaired two-tailed Student’s T-test (H) and two-way analysis of variance (ANOVA) with Bonferroni’s multiple comparisons posttest (I).

**Fig. S6.**
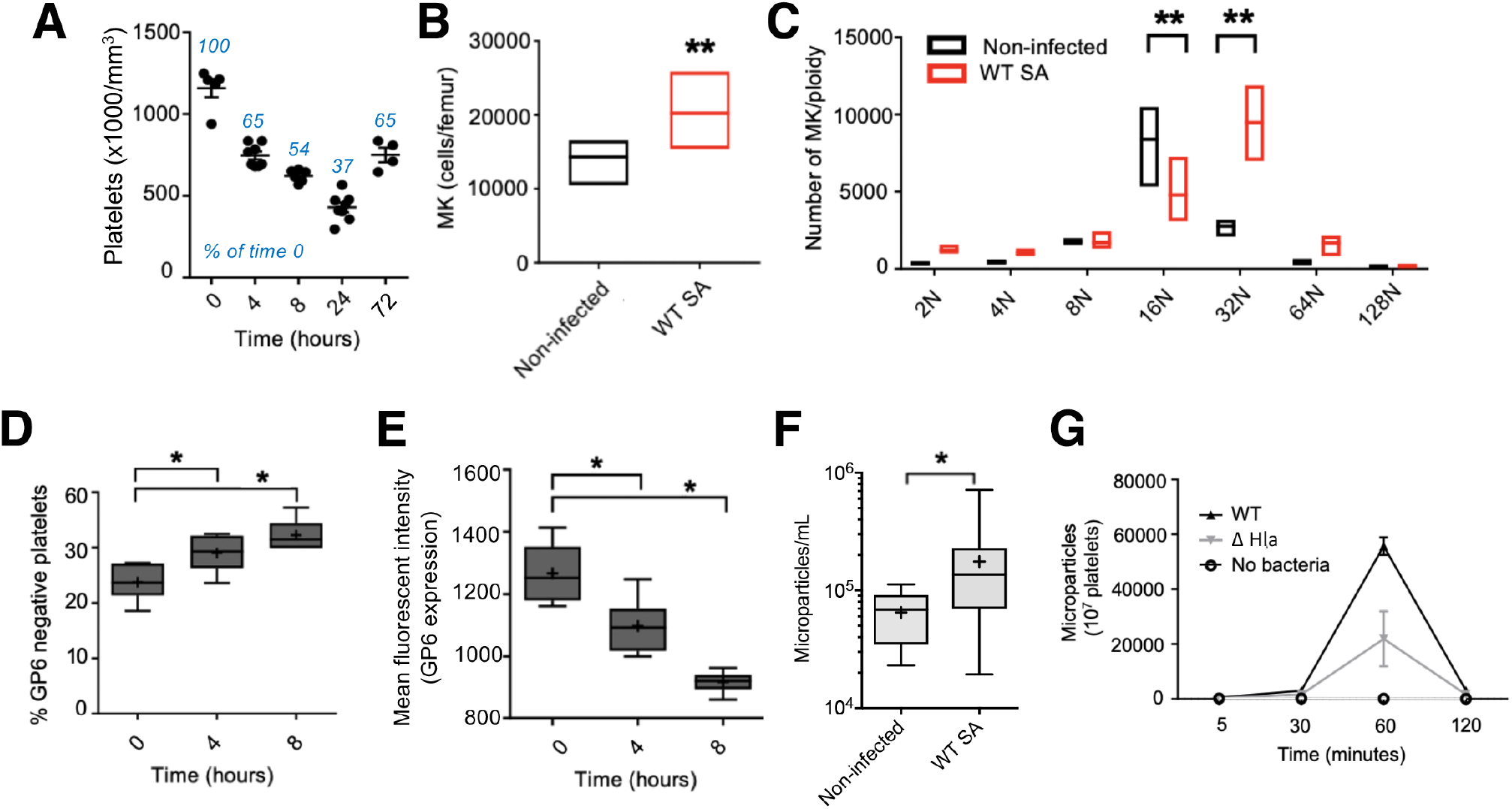
Effects of SA challenge on platelets and thrombopoiesis *in vivo*. (A) Platelet counts after inducing systemic SA infection (5 × 10^7^ CFU SA i.v.). (B and C) Bone marrow cells were harvested after 72 h of SA systemic infection and analyzed for MK count (B) and ploidy distribution (C). (D and E) Circulating platelets were analyzed for % GP6 negative platelets (D) and surface GP6 expression (E) by flow cytometry after inducing systemic SA infection (5 × 10^7^ CFU SA i.v.). (F) Platelet microparticulation was measured *in vivo.* Mice were injected 1 × 10^8^ SA i.v. and after 4 h, blood was collected and analyzed for platelet microparticles by flow cytometry. (G) Human platelets were infected with SA *in vitro.* Platelet microparticles in the samples were analyzed at each time point by flow cytometry.

**Fig. S7.**
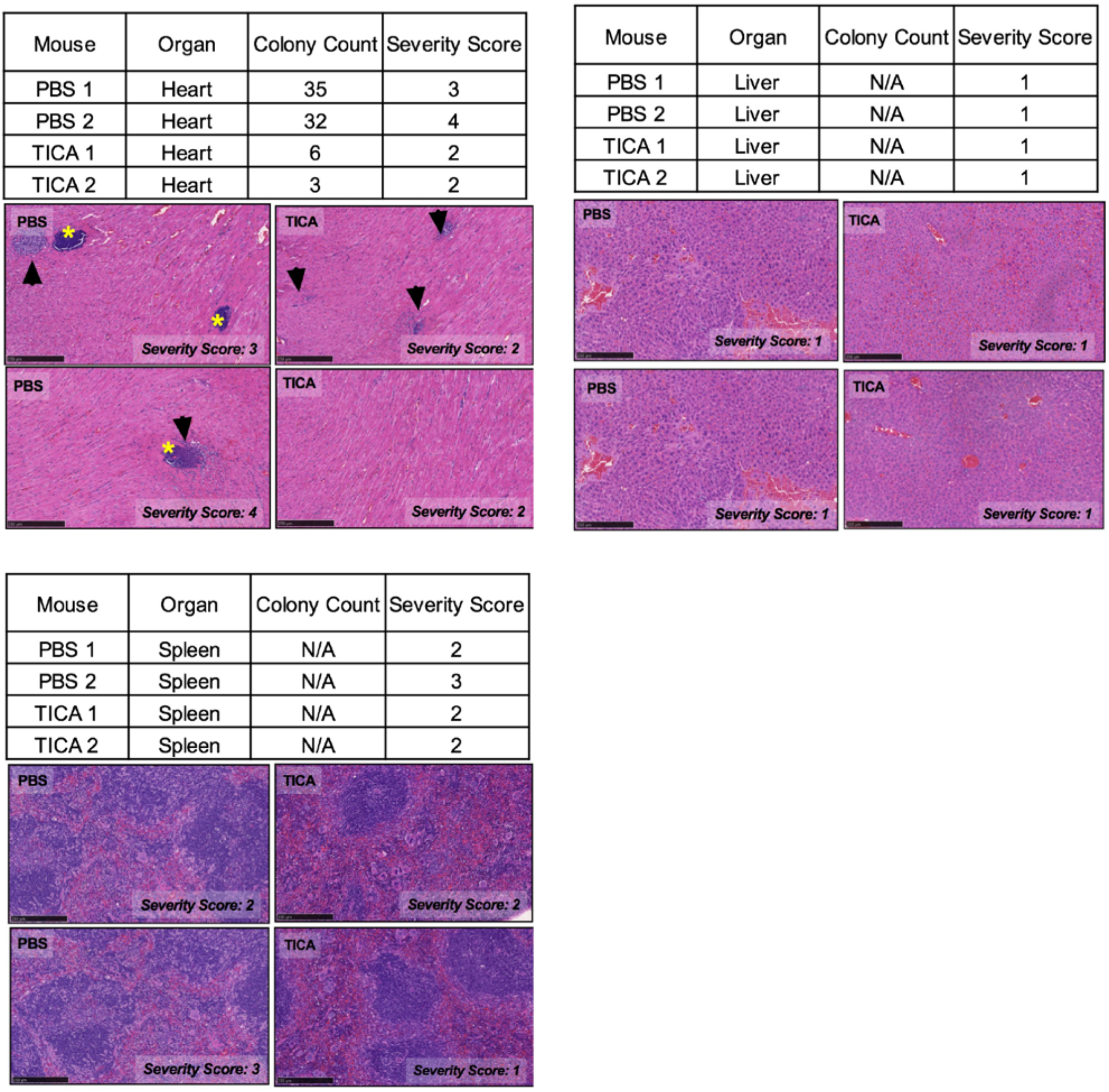
Organ pathology in murine SA infection with or without ticagrelor treatment. (**A**) Hematoxylin and eosin stain (H&E) of representative histological heart, liver, and spleen sections from mice pre-treated with vehicle or 4 mg/kg ticagrelor 12 h prior to SA infection and q 12 h thereafter for 72 h; (*n =* 2). Yellow stars denote dense bacterial colonies. In the Ticagrelor-treated mice, the bacterial colonies were smaller, less frequent, and often surrounded by an inflammatory infiltrate (black arrow) comprising neutrophils with fewer macrophages. Images are representative of at least two independent experiments.

**Fig. S8.**
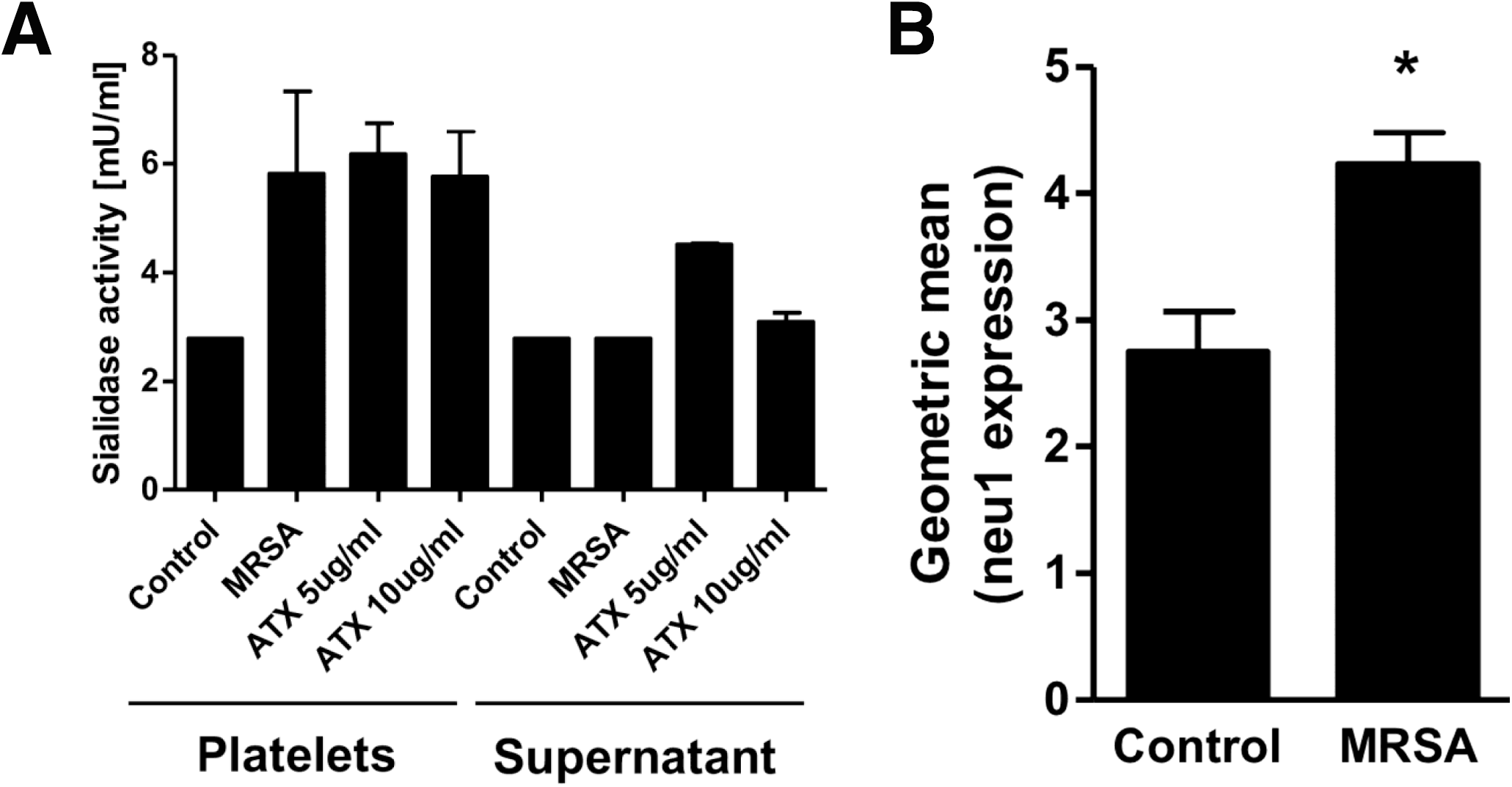
Neu1 is predominantly detected on the platelet cell surface and is induced by SA exposure. (**A**) Human platelet-rich plasma was incubated with 2 × 10^7^ CFU SA or with 5 or 10 μg/ml recombinant α-toxin (ATX). Following 30 min incubation at 37°C, platelets and plasma were separated by centrifugation. Both fractions were assayed for sialidase activity using the Amplyx Red Assay (Fisher Scientific) per manufacturer’s instructions, using DANA as a background control. A standard curve was established using *Arthrobacter ureafaciens* sialidase (AUS), with a starting concentration of 500 mU/mL serially diluted down to 0.98 mU/mL. (B) Human platelets were incubated with WT SA at MOI = 1 for 30 min rotating at 37°C, then incubated with anti-Neu1 antibody, followed by addition of AlexaFluor 488 goat anti-rabbit IgG. Samples were washed and the surface expression of Neu1 analyzed by flow cytometry. Data are shown as mean ± SEM and are representative of at least three independent experiments. Statistical significance determined by unpaired Two-tailed Student’s t test. \**P* < 0.05

**Fig. S9.**
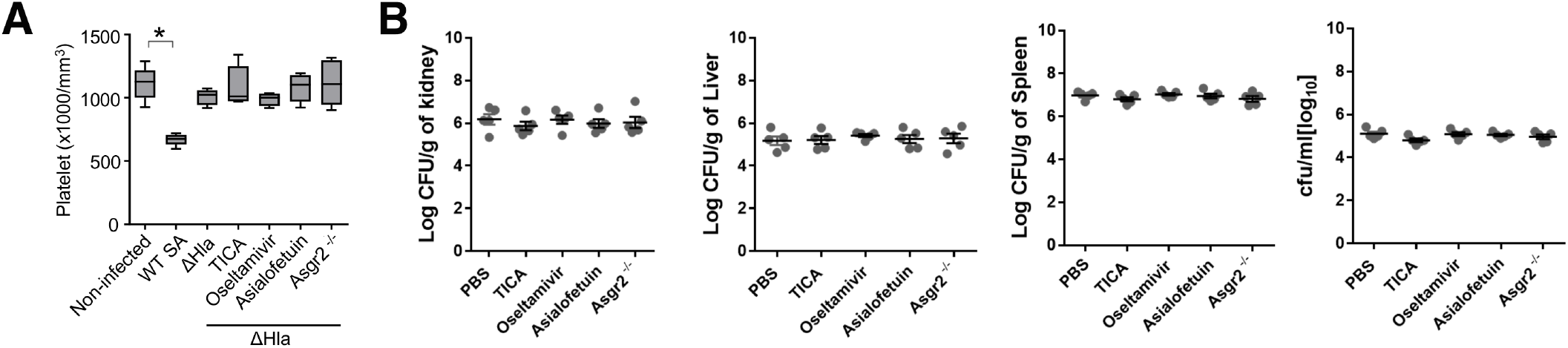
SA α-toxin deletion mutant phenocopies protective effects of TICA, oseltamivir and AMR loss/inhibition. **(A)** WT C57/Bl6 mice or Asgr2^−/−^ mice were either challenged with 8 × 10^8^ WT SA or its isogenic ΔHla mutant. C57/BL6 mice infected with ΔHla were subsequently treated with TICA, oseltamivir, asialofetuin, or PBS control. Blood was harvested 4 h post infection and (A) platelets or **(B)** bacterial colony forming units (CFU) enumerated. Statistical significance determined by unpaired Two-tailed Student’s T-test. For floating bar graphs, - denotes the median, whiskers represent min. to max, and floating box represents the 25th to 75th percentile. Unless otherwise stated, *P < 0.05. PBS, phosphate buffered saline; ns, not significant; WT, wild-type; TICA, ticagrelor.

## Notes

### Competing Interest Statement

The authors have declared no competing interest.

